# Optimization of the precursor supply for an enhanced FK-506 production in *Streptomyces tsukubaensis*

**DOI:** 10.1101/2022.03.16.484622

**Authors:** Susann Schulz, Christoph Schall, Thilo Stehle, Christian Breitmeyer, Sergii Krysenko, Agnieszka Bera, Wolfgang Wohlleben

**Affiliations:** Department of Microbiology and Biotechnology, Interfaculty Institute of Microbiology and Infection Medicine Tübingen (IMIT), University of Tübingen, Auf der Morgenstelle 28, 72076 Tübingen, Germany; Cluster of Excellence ‘Controlling Microbes to Fight Infections’, University of Tübingen, Auf der Morgenstelle 28, 72076 Tübingen, Germany; Novartis AG, Schaffhauserstrasse 101, 4332 Stein, Switzerland; Interfaculty Institute of Biochemistry, University of Tübingen, Auf der Morgenstelle 34, 72076 Tübingen, Germany; Wörwag Pharma GmbH & Co. KG, Flugfeld-Allee 24, 71034 Böblingen, Germany

**Keywords:** pipecolic acid, FK-506 production, tacrolimus, lysine cyclodeaminase, *S. tsukubaensis*

## Abstract

Tacrolimus (FK-506) is a macrolide widely used as immunosuppressant to prevent transplant rejection. Synthetic production of FK-506 is not efficient and costly, whereas the biosynthesis of FK-506 is complex and the level produced by the wild type strain, *Streptomyces tsukubaensis*, is very low. We therefore engineered FK-506 biosynthesis and the supply of the precursor L-lysine to generate strains with improved FK-506 yield. To increase FK-506 production, first the intracellular supply of the essential precursor lysine was improved in the native host *S. tsukubaensis* by engineering the lysine biosynthetic pathway. Therefore, a feedback deregulated aspartate kinase Ask_*St*_* of *S. tsukubaensis* was generated by site directed mutagenesis. Whereas overexpression of Ask_*St*_* resulted only in a 17% increase in FK-506 yield, heterologous overexpression of a feedback deregulated Ask_*Cg*_* from *Corynebacterium glutamicum* was proven to be more efficient. Combined overexpression of Ask_*Cg*_* and DapA_*St*,_ showed a strong enhancement of the intracellular lysine pool following increase in the yield by approximately 73% compared to the wild type. Lysine is coverted into the FK-506 building block pipecolate by the lysine cyclodeaminase FkbL. Construction of a Δ*fkbL* mutant led to a complete abolishment of the FK-506 production, confirming the indispensability of this enzyme for FK-506 production. Chemical complementation of the Δ*fkbL* mutant by feeding pipecolic acid and genetic complementation with *fkbL* as well as with other lysine cyclodeaminase genes (*pip*_*Af*_, *pipA*_*St*_, originating from *Actinoplanes friuliensis* and *Streptomyces pristinaespiralis*, respectively) completely restored FK-506 production. Subsequently, FK-506 production was enchanced by heterologous overexpression of Pip_*Af*_ and PipA_*Sp*_ in *S. tsukubaensis*. This resulted in a yield increase by 65% compared to the WT in the presence of Pip_*Af*_ from *A. friuliensis*. For further rational yield improvement, the crystal structure of Pip_*Af*_ from *A. friuliensis* was determined at 1.3 Å resolution with the cofactor NADH bound and at 1.4 Å with its substrate lysine. Based on the structure the Ile91 residue was replaced by Val91 in Pip_*Af*_, which resulted in an overall increase of FK-506 production by approx. 100% compared to the WT.

## Introduction

Tacrolimus (FK-506) is a 23-membered macrocyclic polyketide that has firstly been isolated from *Streptomyces tsukubaensis* in 1984 (Kino et al., 1987). It exhibits a strong immunosuppressive activity by blocking the calcineurin phosphatase leading to a reduced T-cell proliferation due to the diminished level of interleukin-2, an essential growth factor for activated T-cells (Schreiber &Crabtree, 1992). Although sharing a similar mode of action, FK-506 is much more potent than the well-established immunosuppressant cyclosporin (Jiang &Kobayashi, 1999; Liu et al., 1991), hence arising great pharmaceutical interest (Haddad et al., 2006, Muduma et al., 2016). Nowadays FK-506 is chosen as a first line drug in various clinical areas of application e.g. after organ transplantation or for treatment of inflammatory skin diseases and eczema leading to an emerging commercial and scientific interest (Staatz &Tett, 2004, Lin, 2010). The biosynthetic pathway of FK-506 has been entirely elucidated (Motamedi et al., 1998) and the whole genome sequencing of the FK-506 producer *S. tsukubaensis* NRRL 18488 has been completed (Blazic et al., 2012). Basically, the biosynthesis of the core polyketide of FK-506 is processed by three different polyketide synthases (PKS) FkbA, FkbB and FkbC, catalyzing the condensation of an unusual starter unit derived from the shikimic acid pathway (4,5-dihydroxycyclo-1-enecarboxylic acid (DHCHC)) with ten extender units (two malonyl-CoA, five methylmalonyl-CoA and two methoxymalonyl-CoA). In a following step the non-ribosomal peptide synthetase (NRPS) FkbP is incorporating the sole peptide moiety pipecolic acid, which is derived from L-lysine, into the polyketide chain. It closes the ring structure in a final cyclization step (Andexer et al., 2011, Motamedi et al., 1997, Motamedi &Shafiee, 1998, Mo et al., 2011, Goranovic et al., 2010) before the immature macrolactone is further processed by post-PKS tailoring enzymes resulting in the final compound FK-506 (Motamedi et al., 1996, Chen et al., 2013) (Fig. 1). Pipecolic acid, a non-proteinogenic amino acid, is a key intermediate in the synthesis of a large number of drugs, e.g. pristinamycin (Mast et al., 2011), friulimicin (Müller et al., 2007), meridamycin (Jiang et al., 2011), rapamycin (Gatto et al., 2006), tacrolimus (Turlo et al., 2012), nocardiospin (Bis et al., 2015), tubulysin B (Steinmetz et al., 2004) and others. In fact, pipecolic acid is often even crucial for the bioactivity of secondary metabolites (Min, 2006).

**Fig. 1.**
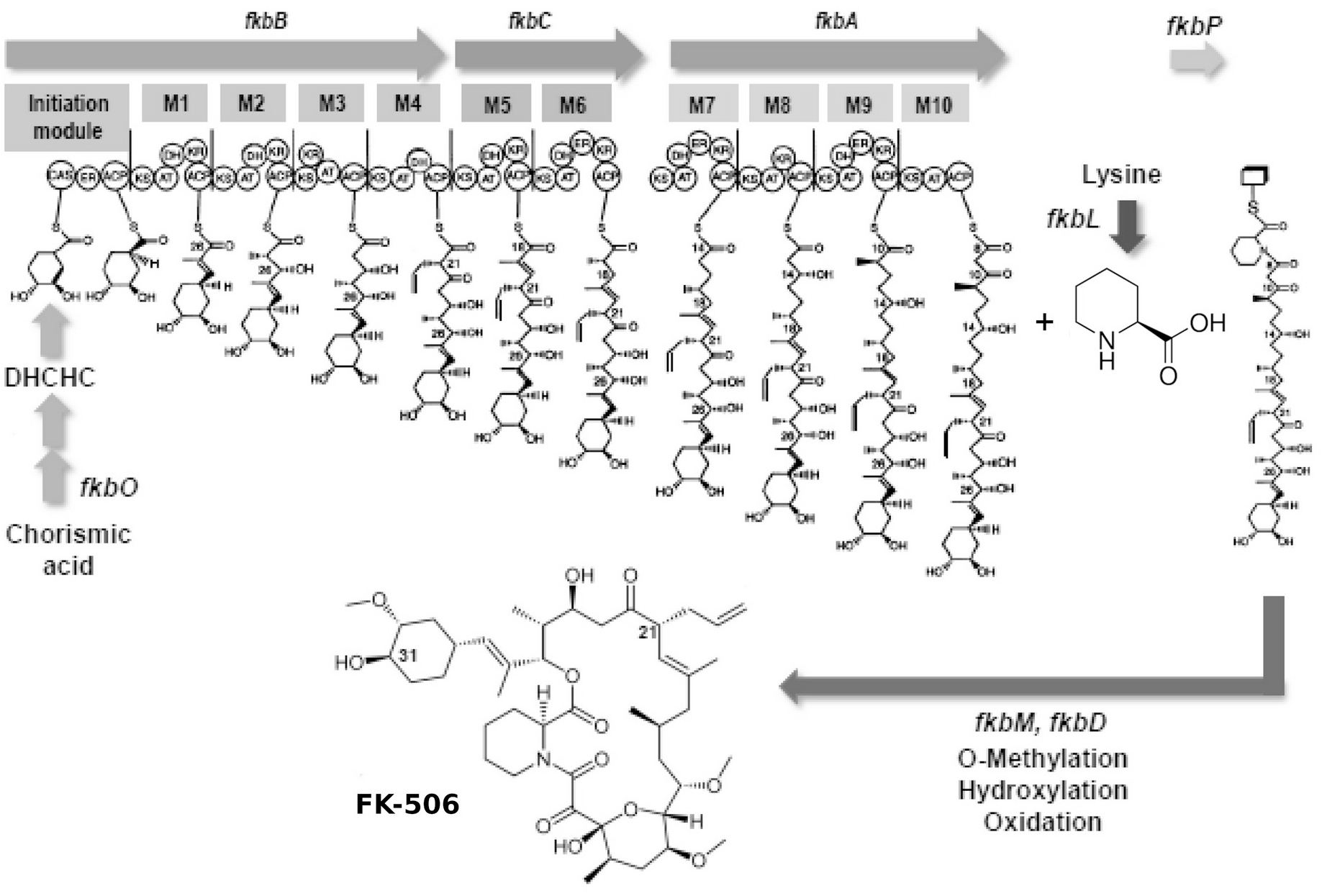
Schematic representation of the FK.506 biosynthesis in *S. tsukubaensis* (modified after Ordóñez-Robles et al., 2018). Arrows in the upper part: three PKS genes (*fkbA, fkbB, fkbC*) and the NRPS gene (*fkbP*) of the biosynthetic cluster. Boxes M1 to M10: Modules of the PKSs; circles are domains in the modules: ACP, acyl carrier protein; AT, acyltransferase; ER, enoyl reductase; CAS, CoA synthetase; KR, 3-oxoacyl (ACP) reductase; DH, 3-oxoacyl thioester dehydratase; KS, 3-oxoacyl (ACP) synthase. DHCHC: (4R, 5R)-4,5-dihydroxycyclohex-1-enecarboxylic acid.

Lately a lot of efforts have been invested in optimizing the biotechnological process of FK-506 fermentation by employing different strategies ranging from classical strain improvement methods to more purposeful metabolic profiling approaches (Yoon &Choi, 1997; Menglei et al., 2013; Ban et al., 2016). Since the major bottleneck for a high FK-506 yield may be the limited supply of building blocks, plenty of the scientific work on the FK-506 production improvement was dedicated to the optimization of increased levels of precursors. Most strategies focused on specific supplementation of the production media with essential precursors. Others directly applied genetic approaches for the targeted engineering of biosynthetic pathways which deliver FK-506 building blocks (Turło et al., 2012; Chen et al., 2012; Huang et al., 2013; Huang et al., 2013). Particularly, an effect of L-lysine on the production of FK-506 as well as other antibiotics, like ß-lactams, has already been shown in previous feeding studies (Mendelovitz & Aharonowitz; 1982; Mendelovitz & Aharonowitz; 1983; Martínez-Castro et al., 2013; Ying et al., 2017).

In last years, an efficient strategy for heterologous *de novo* biosynthesis of the FK-506 intermediate L-pipecolic acid was presented in recombinant *E. coli* cells (Ying et al., 2017). In order to enhance the metabolic pathway from L-lysine to L-pipecolic acid, *pipA*, a lysine cyclodeaminase gene from *S. pristinaespiralis* was introduced. Additional plasmid-based overexpression of *dapA, lysC*, and *lysA* under the control of the strong *trc* promoter and rebalancing of the intracellular pyridine nucleotide concentration increased the pipecolic acid production (Ying et al., 2017). In this work a new optimization strategy that interconnects the enhancement of the precursor supply for the primary pathway of the L-lysine biogenesis with the secondary pathway of pipecolic acid formation in the native host *S. tsukubaensis* is presented. Among the few routes that have evolved to generate pipecolic acid, the lysine cyclic deamination pathway was mainly found in actinobacteria. The FK-506 gene cluster as well as the pristinamycin and friulimicin gene clusters comprise a gene encoding a lysine cyclodeaminase (*fkbL, pipA*_*St*_ and *pip*_*Af*_, respectively). The lysine cyclodeaminase is involved in one of the late steps of the FK-506 biosynthesis and is catalyzing the direct formation of pipecolate from L-lysine (Gatto et al., 2006).This type of enzyme was biochemically characterized and the crystal structure of PipA from *S. pristinaespiralis* has been solved recently (Ying &Chen, 2016). Combining the information deduced from the pipecolic acid biogenesis from different microorganisms and from the deposited crystal structure of the lysine cyclodeaminase PipA_*Sp*_ as well as in this work determined structure of Pip_*Af*_, a unique approach for the enhancement of FK-506 production in S. *tsukubaensis* had been achieved.

## Materials and methods

### Bacterial strains and plasmids

*S. tsukubaensis* NRRL 18488 from the NRRL Culture Collection of the Agricultural Research Service (USA). All mutants are derivatives of *S. tsukubaensis* NRRL 18488. *Escherichia coli* Novablue cells from Novagen were used for standard cloning procedures, while *E. coli* Rosetta DE2 cells served as hosts for protein expression experiments. For plasmid transfer intergeneric conjugation between the non-methylating *E. coli* ET12567/pUZ8002 strain and *S. tsukubaensis* was used (Kieser et al., 2000). For overexpression experiments the integrative pRM4 vector containing the strong constitutive *ermE* promoter was used (Menges et al., 2007).

For heterologous expression experiments concerning the lysine cyclodeaminases, the *pipA* gene was amplified from genomic DNA from *Streptomyces pristinaespiralis* Pr 11 (Aventis Pharma) (Mast et al., 2011). As template for the amplification of the *pip* gene from *Actinoplanes friuliensis* the plasmid pRF37 from C. Müller *et. al*. 2007 was used (Müller et al., 2007). *S. tsukubaensis* NRRL 18488 was used as a template for the amplification of the *fkbL, fkbP* and *dapA*.

The genomic DNA for the amplification of the feedback inhibition deregulated aspartate kinase gene *lysC*_*Cg*_* from *Corynebacterium glutamicum* DM1730 (Seibold et al., 2006) was kindly provided by Jörn Kalinowski (University of Bielefeld, Germany).

All strains and plasmids used in this study are listed in the Suppl. Table 2. All oligonucleotides used in this work are listen in the Suppl. Table 3.

### Media and culture conditions

Spores and mycelia preparations were obtained from ISP4 (Difco, Sparks, MD, USA) agar plates. FK-506 production by *S. tsukubaensis* NRRL 18488 was analyzed in the liquid media MG optimized by Martínez-Castro *et al*. 2013. FK-506 production was also analyzed in R2YE medium (Kieser et al., 2000) modified by Temuujin *et al*., 2011. The composition of MGm medium was as follows: 50 g/l starch (Difco), 8.83 g/l glutamic acid, 2.5 mM KH_2_PO_4_/K_2_HPO_4_, 0.2 g/l MgSO_4_·7H_2_O, 1 mg/l CaCl_2_, 1 mg/l NaCl, 9 mg/l FeSO_4_ ·7H_2_O, 21 g/l MOPS, and 0.45 ml/l 10× trace elements, adjusted to pH 6.5. The composition of R2YE medium was as follows: 103 g/l saccharose, 0.25 g/l K_2_SO_4_, 10.12 g/l MgCl_2_*6H_2_O, 10 g/l glucose, 0,1 g/l casamino acids, 5 g/l yeast extract, 10 ml/l 0,5% K_2_HPO_4_, 80 ml/l 3,68% CaCl_2_*2H_2_O, 15 ml/l 20% L-prolin, 100 ml/l 5,73% TES, 2 ml/l trace elements, pH 7,2. For 10 ml solution (100×) of trace elements, the components are: 39.0 mg CuSO_4_ ·5H_2_O, 5.7 mg H_3_BO_3_, 3.7 mg (NH_4_)_6_Mo_7_O_24_·4H_2_O, 6.1 mg MnSO_4_·H_2_O, and 895.0 mg ZnSO_4_·7H_2_O. Fermentation was performed using a two-stage culture system. For the seed culture a mixture of 1:1 modified YEME (3 g/l yeast extract, 5 g/l Bacto peptone, 3 g/l malt extract, 10 g/l glucose and 340 g/l sucrose) and Tryptic Soy Broth (Difco) was inoculated with spores or mycelia from an ISP4 plate and cultivated for at least 3 days at 28°C and 220 rpm. For the main culture 100 ml of MGm medium were inoculated with the seed culture and further incubated at the same condition for 6 days.

### Analysis of growth and FK-506 production

The productivity of an *S. tsukubaensis* strain was usually determined as production of FK-506 per g cell dry weight. During the fermentation time of 6 days all 24h 1 ml of bacterial culture was gathered, centrifuged, washed twice with destilated water, and dried by lyophillization. For FK-506 determination, the broth samples were mixed with equal volume of ethylacetate (1:1), stirred for 10 min and subsequently centrifuged. The organic phase was analysed via an HP1090 M HPLC system equipped with a diode array detector and a thermostatic autosampler. A Zorbax Bonus RP column, 3 × 150, 5 μm (Agilent Technologies, Santa Clara, USA) constituted the stationary phase. The mobile phase system was applied with 0.1% phosphoric acid and 0.2% triethylamine as eluent A and acetonitrile with 1% tetrahydrofurane as eluent B. The flow rate was 850 μL/min, the column temperature was set to 60°C. The absorbance was monitored at a wavelength of 210 nm. Data sets were processed with the help of the Chemstation LC 3D, Rev. A.08.03 software from Agilent Technologies. Standards of pure FK-506 (Antibioticos SA) and ascomycin (Sigma-Aldrich Chemie GmbH, Munich, Germany) were used as controls.

### Homology protein modelling and analysis of protein 3D structures

A 3D protein model of a target sequence was generated in comparative modelling by extrapolating experimental information from an evolutionary related protein structure that serves as a template using SWISS-MODEL (Waterhouse et al., 2018). Template search was based on the target sequence, which served as a query to search for evolutionary related amino acid sequences by BLASTp (Altschul et al., 1997) and for protein structures against the SWISS-MODEL template library. Templates were ranked according to expected quality of the resulting models and used to automatically generate 3D protein models by loop modelling. The quality of obtained models was estimated based on expected model accuracy, the QMEAN scoring function (Studer et al., 2020) as well as based on the Ramachandran plot (Ramachandran et al., 1963). 3D models of proteins were analyzed using the Swiss-PdbViewer software (Guex et al., 2009).

### Expression and purification of the His-tagged Pip protein

The *pip*_*Af*_ gene was amplified using the above mentioned template from Müller *et. al. 2007*. It was cloned in the expression vector pET30 Ek/lic containing an N-terminal His-tag with the help of the ligation independent Ek/lic system according to the protocol proposed by Novagen, UK. *E*.*coli* Rosetta DE2 was transformed with the resulting construct pET30/pip. 200 ml of Luria Bertami broth (LB) (Sambrook &Russell, 2006) with kanamycin (50 µg/ml) and chloramphenicol (34 µg/ml) was inoculated with 2 ml of an overnight culture and incubated at 37°C on a rotary shaker (180 rpm). Expression was induced with 1mM IPTG after the culture broth has reached an OD_578_ of 0.4. The induced culture was further incubated for 18h. The cells were harvested by centrifugation, resuspended in lysis buffer (20 mM Tris, 100 mM NaCl, 20 mM imidazol, protease inhibitor (Roche, Mannheim, Germany)) and lysed by 3-4 passages through Emulsiflex B15 (Avestin, Ottawa, Canada). Protein purification was either carried out by gravity flow Ni NTA Superflow columns (Iba) or FPLC using a His-TrapFF 1ml column (GE Healthcare, Munich, Germany). Concentrated protein eluates were stored at 4°C.

### Construction and expression of Pip* variants

Mutated Pip* variants were generated by site-directed mutagenesis. The procedure was based on the Stratagene quickchange protocol, which was optimized by Zheng *et al* i (2004). For each mutation specific primers were designed. Mutations were introduced into the pET30/*pip* expression plasmid using PCR. Non-mutated template DNA was digested using the restriction enzyme D*pn*I. Subsequently, the competent *E. coli* Novablue strain was transformed. Plasmid-carrying clones were identified by selection for kanamycin and verified by PCR and sequencing.

### Crystallization and structure determination of Pip

The obtained Pip_*Af*_ was concentrated by Amicon Ultra spin concentrators and further purified by gel filtration on a Superdex 200 run with 150 mM NaCl and 25 mM HEPES pH 7.4. The main peak corresponding to a dimer was pooled and concentrated up to absorption at 280 nm of 2.5 equal to a protein concentration of 9.5 mg/ml. The absorption at 340 nm thereof varied from 0.4 to 0.8 depending on the batch. For crystallization the concentrate was supplemented with 1 mM NAD^+^ and diluted to 7.3 mg/ml. The same solution was also used for co-crystallization with the substrate but contained additionally lysine to a final concentration of 140 mM. The crystallization drops contained 450 nl of protein and 300 nl of crystallization solution and were set up in sitting drop microtiterplates with a Tecan Freedom Evo pipetting robot. First crystals grew after few hours at 4°C in several conditions. Lysine-Cocrystallization was optimized with JCSG + suite screen (Qiagen, Hilden, Germany) condition 42 (20% PEG8000, 0.2 M MgCl_2_, 0.1 M Tris pH 8.5) and 10% glycerol added before freezing. From the setups without lysine crystals were obtained from condition 36 of Morpheus screen (Molecular dimensions) which were of sufficient quality.

The crystals were mounted in loops and flash-frozen in liquid nitrogen for storage and measured at the Swiss Light Source (SLS, Villingen, Germany). Data reduction was carried out with the XDS package. For phase determination with PHASER the OCD structures were truncated and used as search model. The structure was built in interactive cycles with coot and for refinement; simulated annealing was included in the very first refinement run. The representation of the solved protein structure was carried out using the free graphics 3D software PyMOL (The PyMOL Molecular Graphics System, Version 2.0 Schrödinger, LLC).

### Thin layer chromatography for qualitative detection of pipecolic acid

Overnight cultures of expression strains *E. coli* Rosetta 2 (pET/*pip*_*Af*_) and *E. coli* Rosetta 2 (pET/*pip*_*Af*_*\**) and the control *E. coli* Rosetta 2 were incubated overnight at 37°C in 10 ml LB medium with the corresponding kanamycin and chloramphenicol concentrations. Subsequently, 100 ml of LB_KM/CM_ of the main culture was supplemented with 1 µL of the corresponding pre-culture and incubated up to an OD_578_ of 0.6 at 37°C. After reaching the OD_578_ of 0.6, the expression of the Pip_*Af*_/Pip_*Af*_* proteins was induced by adding 1mM IPTG with the subsequent incubation at 28°C for further 20 h. At the next day, the main cultures were supplemented with 0.5 ml of the glycerol stock solution (50%) as well as 0.5 ml lysine stock solution (50%) and incubated for 6 h at 32°C. Afterwards, the cultures were centrifuged for 10 min at 5000 rpm. To determine the formation of pipecolic acid, 5 µL of supernatant from each culture was spotted at a silica gel plate. The mobile phase, which consisted of a mixture of 3:1:1 N-butanol:acetic acid:water, was used for the separation of substances. As controls the pure lysine and pipecolic acid (5 µg each) were used. After the running front had reached a height of approx. 10 cm, the silica gel plate was stained with 0.5% ninhydrin solution and analyzed. Quantification of pipecolic acid on TLC was performed using the open-source software ImageJ: percentage of each peak was related to the total area of all peaks and represents the relative amount of the pipecolate based on digital imaging.

## Results and Discussion

### Strategies for FK-506 production optimization

Analysis of randomized controlled trials showed FK-506 to be superior to ciclosporin in terms of patient mortality and hypertension (Muduma et al., 2016). However, the biotechnological production is limited due to the low titers of FK-506 produced by the wild type strains. While some groups focused on classical feeding strategies by supplementing relevant precursors and media optimization (Menglei et al., 2013; Martínez-Castro et al., 2013; Ban et al., 2016), others tried to solve the problem by genetic manipulation targeting metabolic pathways, e.g. via overexpression of FK-506 biosynthetic genes (Huang et al., 2013; Fu et al., 2016; Ban et al., 2016; Ordóñez-Robles et al., 2018). In order to optimize the precursor flux we aimed to combine optimization of specific natural product synthesis in the endogenous system as well as the synthetic biology approach with genes from bacteria that are used for the biotechnological production of amino acids.

### Improvement of the lysine precursor supply in *Streptomyces tsukubaensis*

L-lysine is an essential amino acid, used in nutrition, as supplementary and nutraceutical, which production has been established in *C. glutamicum*. Also, lysine is an important product that serves as a precursor for L-pipecolinic acid (piperidine-2 carboxylic acid), which is a non-proteinogenic amino acid widely used in all phylogenetic domains of life and which is a building block of biologically active substances such as streptogramin pristinamycin (Mast et al., 2011) as well as macrocyclic immunomodulators such as rapamycin and FK-506 (Turlo et al., 2012). Enhancing the L-lysine precursor pool should result in a positive effect on secondary metabolite production. To proof this hypothesis classical media supplementation experiments with various amino acids have been performed showing that particularly exogenous L-lysine feeding lead to a significant boost of rapamycin (Cheng et al., 1995) respectively FK-506 production (Huang et al., 2013; Martínez-Castro et al., 2013), consequently making the L-lysine primary biosynthesis pathway a putative target for genetic engineering. For instance, enrichment of the semi-defined medium for FK-506 production in *S. tsukubaensis* with 2.5 g/l L-lysine enhanced the production by approximately 30% (Martínez-Castro et al., 2013). In order to test, whether additional lysine can further improve the FK-506 yield, we extended this experiment by increasing the exogenous lysine end concentration beyond 2,5 g/l. However, at higher lysine concentration (50 mM) the FK-506 production in the wild type was clearly inhibited (Suppl. Fig. 3)

Therefore, we intended to improve the intracellular L-lysine supply by adjusting the lysine biosynthetic pathway in *S. tsukubaensis* for optimal FK506 production. The biosynthesis of lysine has extensively been studied in microorganisms leading to a biotechnological production in high level producer strains like *C. glutamicum* and *E. coli* (Sahm et al., 1996). In *E. coli*, the combined plasmid-based overexpression of *dapA, lysA* and *lysC* from *E. coli* under the control of the strong *trc* promoter as well as the overexpression of *ddh* from *C. glutamicum* were studied for their role in the lysine biosynthesis to increase the biosynthetic pool of L-lysine (Ying et al., 2017). But this pathway has not specifically been optimized for the production of pipecolate containing metabolites in actinobacteria, taxonomically related to corynebacteria. Studies in *C. glutamicum* provided good understanding of the lysine biosynthesis. This process includes two central optimization steps involving the dihydropicolinate synthase DapA, which is the branching point between lysine and threonine pathway, and the aspartate kinase Ask, which catalyzes the first step in this biosynthetic pathway and is strictly regulated by its end products lysine and threonine (Fig. 2). It has been shown that high-level lysine producers of *C. glutamicum* possess a defect in feedback inhibition of the corresponding aspartate kinases (Garcia et al., 2016). Thus, we hypothesized that central bottlenecks of lysine biogenesis in *S. tsukubaensis* that need to be overcome are the feedback inhibition of the aspartate kinase and the carbon flux towards the lysine branch. We therefore chose the dihydropicolinate synthase (DapA_*St*_) and the aspartate kinase (Ask_*St*_) to be processed to direct the precursor flux towards lysine and hence towards elevated levels of the building block pipecolic acid.

**Fig. 2.**
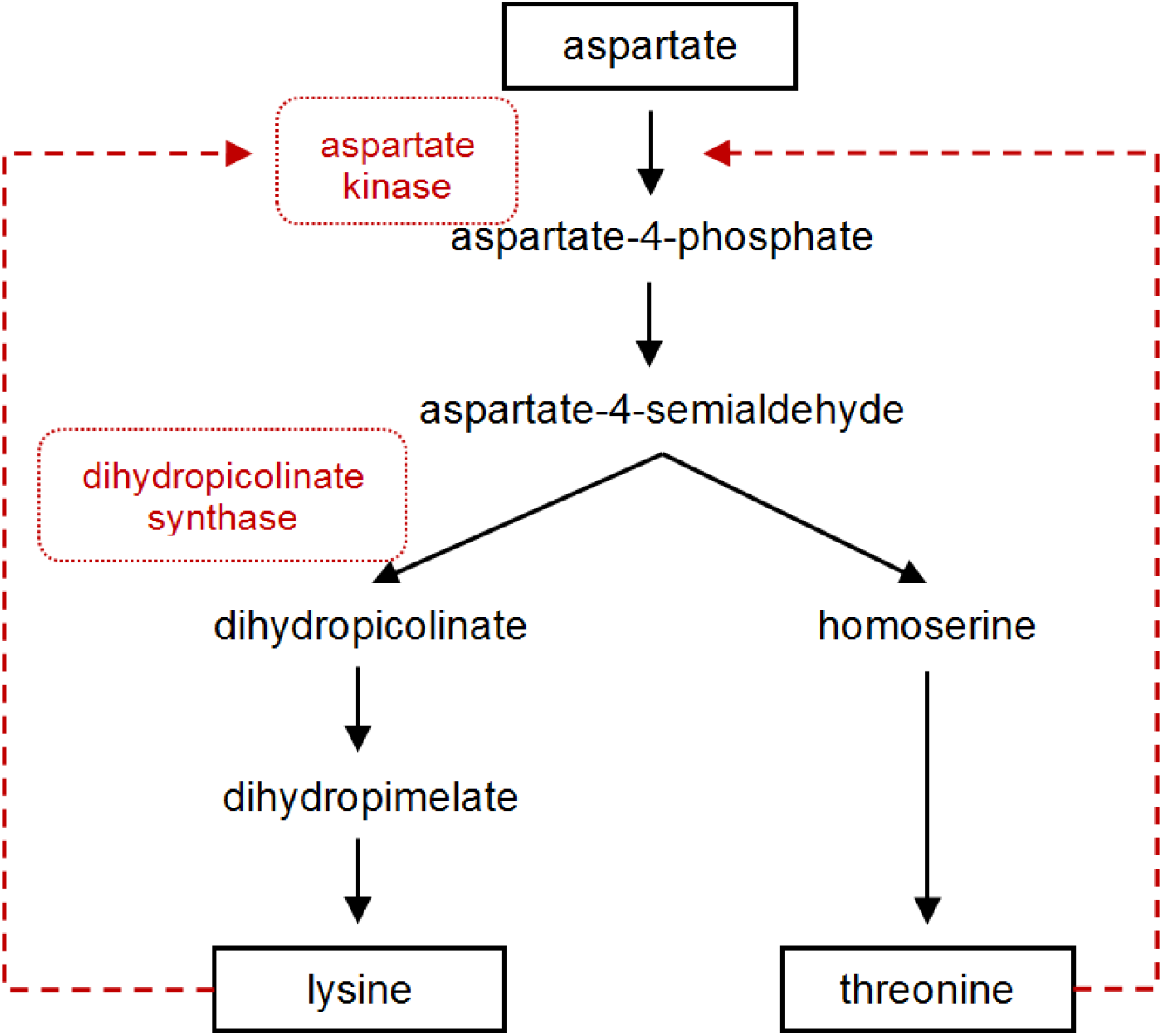
The postulated lysine and threonine biosynthesis pathway in *S. tsukubaensis*. Dashed arrows represent feedback inhibition. Enzymes that were overexpressed in this study are marked in red.

Overexpression of both enzymes in the wild type producer *S. tsukubaensis* NRRL18488 individually did not result in a significant increase of FK-506 yield (Fig. 2). Considering the feedback regulation of the aspartate kinase as the major bottleneck in lysine biosynthesis, it is not surprising that overexpression of the dihydropicolinate synthase (DapA_*St*_) alone does not lead to a profound effect on FK-506 production (∼7%). The overexpression of Ask _*St*_ alone demonstrated no significant effect on FK-506 production (∼8%) (Fig. 3), since this enzyme is subject to strict regulation by a final product inhibition lysine and threonine (Fig. 2).

**Fig. 3.**
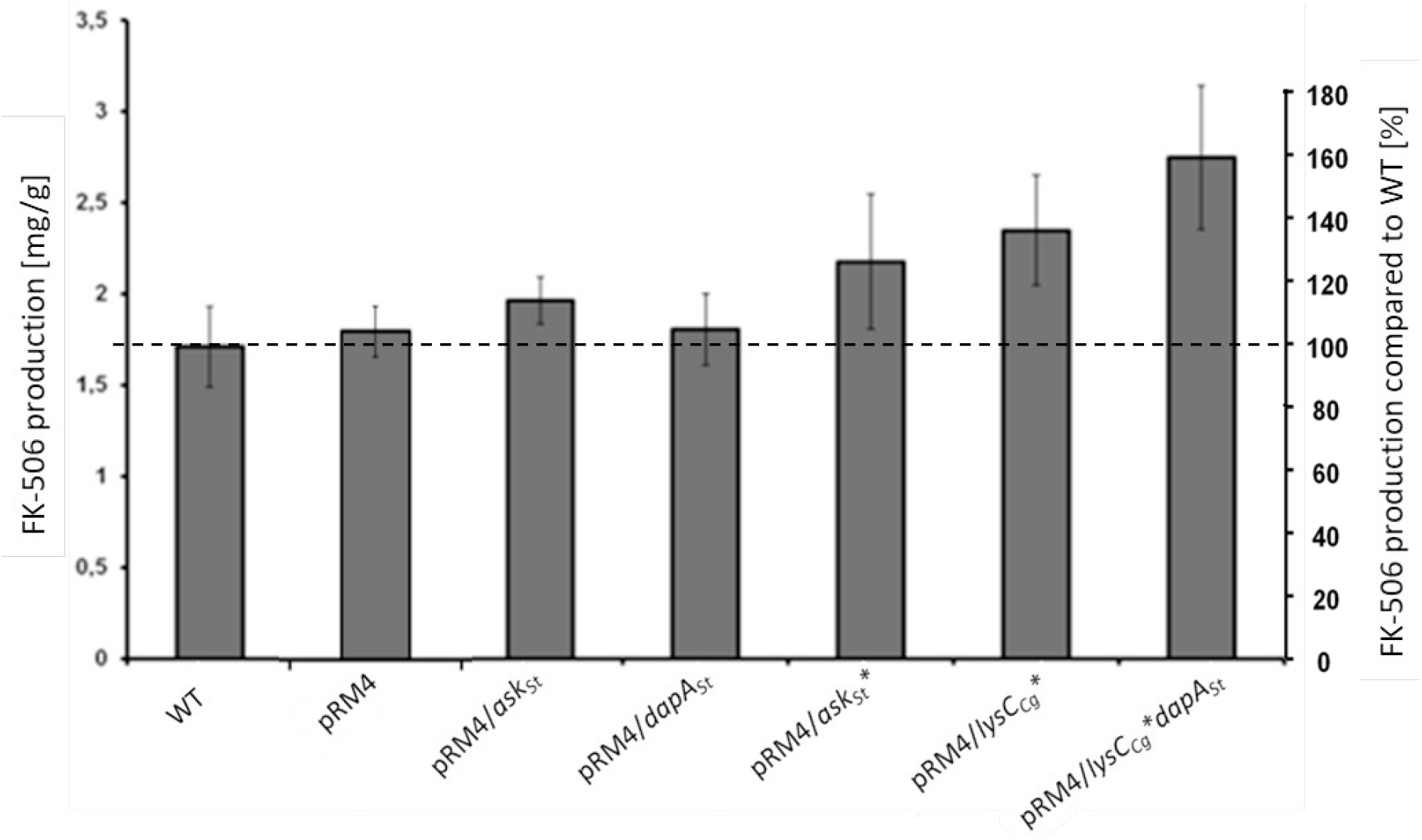
Comparison of the FK-506 production of the wild type and the overexpression strains with optimized lysine supply. X-axis on right: percentage representation of production optimization; as 100% the FK-506 production of the wild type (WT) was set at the respective time. X-axis on left: calculation of the mean of specific FK-506 production over a fermentation period of six days. pRM4 stays for the integrative plasmid without an inserted gene. pRM4/ - integration of an additional copy of the gene on the integrative plasmid pRM4 encoding: *ask*_*St*_ – aspartate kinase; *dapA*_*St*_ - dihydropicolinate synthase; *ask*_*St*_*\** – feedback insensitive aspartate kinase; *lysC*_*Cg*_*\** – deregulated aspartate kinase from *C. glutamicum*.

The primary strategy to relieve this bottleneck was constructing a feedback insensitive aspartate kinase (Ask_*St*_*) using the data of Krämer *et al*. 2006 available for the aspartate kinase from the actinomycete *C. glutamicum*. For this purpose a single amino acid exchange in the wild type Ask (from Ser^301^ to Tyr^301^) was introduced via site-directed mutagenesis. The resulting Ask_*St*_* was overexpressed in *S. tsukubaensis* leading only to a slight enhancement of approximately 17% of FK-506 production (Fig. 3).

Therefore, a second strategy involving the heterologous expression of a gene encoding a deregulated aspartate kinase (Ask_*Cg*_* = LysC_*Cg*_*) (Seibold et al., 2006) from the high-level lysine producer strain *C. glutamicum* ATCC 13032 has been used. This attempt resulted in a significant FK-506 yield improvement of about 46 % (Fig. 3) delivering a more beneficial outcome than supplementation of the production medium with lysine only reaching 30% improvement. Combining overexpression of both enzymes, the deregulated aspartate kinase (LysC_*Cg*_*) together with the dihydrodipicolinate synthase (DapA_*St*_), consistently shifted the carbon flux toward L-lysine formation and resulted in a considerable upgrade of FK-506 production of about 73% (Fig 3). This demonstrates the advantage of genetic engineering in contrast to exogenous feeding approaches. In cephamycin producing *S. clavuligerus* strain a positive correlation between deregulated aspartate kinases and the improvement of antibiotic production was also previously described (Özcengiz et al., 2010).

### Improving the pipecolic acid precursor supply in *Streptomyces tsukubaensis*

Pipecolic acid represents a lysine derived, non-proteinogenic amino acid embedded in the polyketide core structure of the FK506 molecule. In some microorganisms, the biosynthesis of the pipecolic acid has been described to occur via two-step biosynthesis routes (Gupta &Spenser, 1969; He, 2006; Tani et al., 2015). In contrast, during the biosynthesis of rapamycin in *S. hygroscopicus* the conversion of L-lysine to L-pipecolic acid in one step by a lysine cyclodeaminase (LCD) has been shown to be directly catalyzed (Gatto et al., 2006). The one-step pipecolic acid synthesis via the LCD has the advantage to be more suitable for combining it with upstream metabolic processes (Gatto et al., 2006). For example, it was reported that a recombinant *E. coli* strain overexpressing LCD *pipA*_*Sp*_ could produce L-pipecolic acid from L-lysine with a yield increase of nearly 70% compared to the parental strain *E. coli* BL21(DE3) (Ying et al., 2015).

To ensure the supply of the pipecolic acid in *S. tsukubaensis*, the FK506 biosynthetic gene cluster contains the gene *fkbL*. Due to the similarity of FkbL to ornithine cyclodeaminases, the product of *fkbL* is predicted to be a LCD that catalyzes the conversion of L-lysine to pipecolic acid in a deaminative cyclization reaction. The resulting non-proteinogenic amino acid is then activated and attached to the polyketide chain by the NRPS FkbP. The pipecolate moiety plays a crucial role for the biological activity of FK-506 as it is part of the binding motif for the cognate immunophilin (FKBP-12) in T-cells (Andexer et al., 2011).

The essentiality of FkbL for FK-506 production was tested by the construction of an *fkbL* deletion mutant in *S. tsukubaensis* through the replacement of *fkbL* by apramycin resistance gene, which resulted in a tacrolimus null-mutant. Subsequently, the *fkbL* mutant was complemented with *fkbL*. The FK-506 production in the mutant and the recombinant strains grown in the MG production medium was analyzed by HPLC. The loss of FK-506 production in the *ΔfkbL* mutant of *S. tsukubaensis* could also be reconstituted by exogenous addition of pipecolate, which again proved the indispensability of this building block for FK-506 biosynthesis (Suppl. Fig. 1, 2).

In order to test whether overexpression of the essential *fkbL* gene increases FK-506 production, the plasmid pRM4/*fkbL* including the *fkbL* gene under control of the constitutive *ermE\** promoter was introduced into *S. tsukubaensis* WT. The resulting construct revealed a yield increase of 47%. (Fig. 4).

**Fig. 4.**
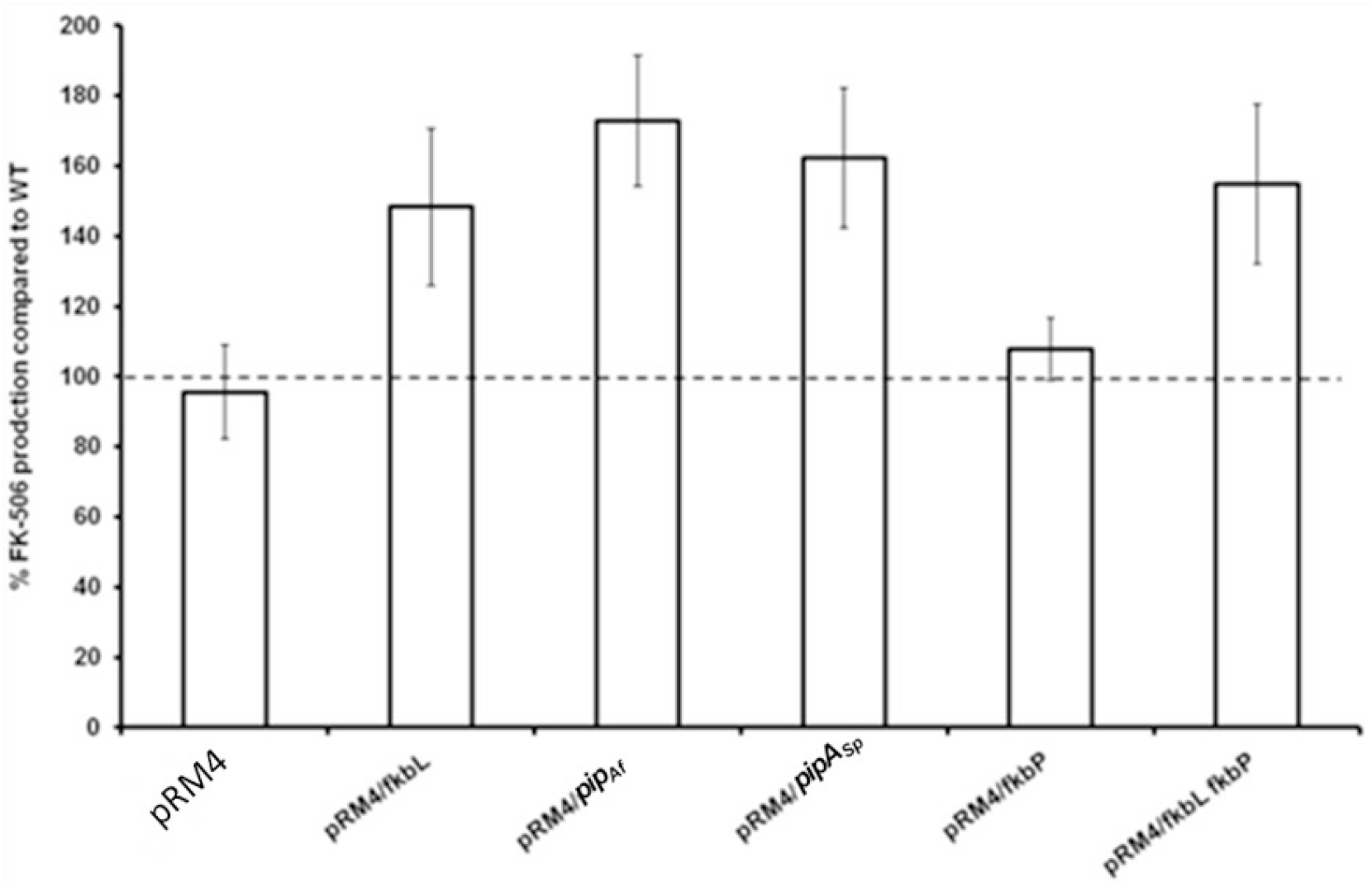
Comparison of the FK-506 production of the wild type and the overexpression strains with improved pipecolic acid supply. pRM4 stays for the integrative plasmide without an incerted gene. PRM4/ - integration of an additional copy of the gene on the integrative plasmide pRM4 encoding: *fkbL* – LCD FkbL from *S. tsukubaensis*; *pip*_*Af*_ - LCD Pip_*Af*_ from *A. friuliensis*; *pipA*_*Sp*_ – LCD PipA_*Sp*_ from *S. pristinaespiralis*; *fkbP* – NRPS FkbP. FK-506 production was measured as an average from three replicates.

Pipecolate is incorporated into the growing FK-506 backbone by the NRPS FkbP, which could constitute a limiting factor for FK-506 production. Therefore, the effect of overexpression of *fkbP* was analyzed. However, only a slight yield improvement (11%) was observed when *fkbP* alone was overexpressed. However, combination of overexpression of both genes *fkbL* and *fkbP* led to a final increase of ∼55% (Fig. 4). These results were acquired from production assays, which were carried out in flask cultivation with the chemically defined medium. Similar observations were reported from fermentation studies with *S. tsukubaensis* D852, another FK-506 producer (Huang et al. 2013). For this study, genes *fkbO, fkbL, fkbP, fkbM* and *fkbD* were introduced into the parental strain (each as a single construct) using the integrative *E. coli–Streptomyces* vector pIB139 containing the *ermE*\* promoter (P*ermE* *) and overexpressed.

Since apparently the production of pipecolate is a highly relevant criteria for efficient FK-506 production, this step of the biosynthesis was analyzed in more detail. Pipecolic acid is a widely distributed constituent of secondary metabolites like for example the streptogramin antibiotic pristinamycin I from *Streptomyces pristinaespiralis* (Cocito, 1979; Mast et al., 2011) and the lipopeptide friulimicin B from *Actinoplanes friuliensis* (Aretz et al., 2000; Müller et al., 2007). In these microorganisms the biosynthesis of the pipecolate building block is also catalyzed by a lysine cyclodeaminase encoded by *pipA*_*Sp*_ in *S. pristinaespiralis* (Mast et al., 2011) and by *pip*_*Af*_ in *A. friuliensis* (Müller et al., 2007). The products of both genes, PipA_*Sr*_ and Pip_*Af*_, show a protein identity of 57% and 50% to the wild type FkbL, respectively. In order to test, whether the heterologous lysine cyclodeaminases can replace the endogenous enzyme, each of the lysine cyclodeaminase genes was expressed heterologously in the *ΔfkbL* mutant under the control of the constitutive *ermE\** promoter. Indeed PipA_*Sp*_ as well as Pip_*Af*_ were able to restore FK-506 production demonstrating that both enzymes are able to form pipecolic acid needed for FK506 biosynthesis (Suppl. Fig. 1, 2). In order to compare the potential of the PipA_*Sp*_ and Pip_*Af*_ for the improvement of FK-506 we overexpressed both *pipA*_*Sp*_ and *pip*_*Af*_ in *S. tsukubaensis* wild type under the control of the constitutive *ermE\** promoter. An improvement of FK-506 production of 56% in case of *pipA*_*Sp*_ and 65% in case of *pip*_*Af*_ overexpression was observed (Fig. 4).

### Identification of structural features of the binding pocket in PipA_*Sp*_ and its comparision with the model structures of Pip_*Af*_ and FkbL

Regarding the significant sequence homology of PipA_*Sp*_ and Pip_*Af*_ to FkbL, it was rather unexpected to see a substantial difference in FK-506 enhancement. To understand the basis for this effects, we aimed to analyze and compare the structures of the three lysine cyclodeaminases PipA_*Sp*_, Pip_*Af*_ and FkbL, to optimize subsequently the effectiveness of the FK-506 production. Resolving of the LCD structure should provide a base for precise enzyme classification, understanding its properties, and future enzyme engineering.

The crystal structure of the LCD PipA_*Sp*_ from *S. pristinaespiralis* was reported and deposited in the protein data base PDB as 5QZJ (Ying &Chen, 2016). Also, the biochemical activity of PipA_*Sp*_ has previously been studied (Gatto et al., 2006, Tsotsou &Barbirato, 2007, Byun et al., 2015). The crystal structures of Pip from *A. friuliensis* as well as FkbL from *S. tsukubaensis* remained unknown so far. In order to compare the structural composition of PipA_*Sp*_, Pip_*Af*_ and FkbL, first homology modelling of Pip_*Af*_ and FkbL was performed. The structures of Pip_*Af*_ and FkbL were generated based on the template with the best query cover and known crystal structure – PipA_*Sp*_.

Although the PipA_*Sp*_ structure has been deposited in the PDB database, its properties and key amino acids in the binding pocket have not been described. PipA_*Sp*_ belongs to the µ-crystallin family and is related to ornithine cyclodeaminases (OCD). It is annotated as lysine decarboxylase WP_037775551.1, alongside with FkbL as WP_006350828.1 and other proteins including AF235504.1 in *S. hygroscopicus* (Wu et al., 2000) and AGP59511.1 in *S. rapamycinicus* (Baranasic et al., 2013). In contrast, Pip_*Af*_ is annotated in the NCBI database as OCD WP_023362365.1.

In order to identify possible key amino acids in the catalytic center of PipA _*Sp*_, we compared it with the structurally solved OCD from *Pseudomonas putida* _*Pp*_OCD (Goodman at al., 2004) with PDB accession code 1×7D, using the Swiss-Model Structure Comparision tool and Swiss PDB-Viewer Magic Fit tool. PipA_*Sp*_ consists, similar to described OCD, of a homodimeric fold whose subunits comprise two functional regions: a substrate-binding domain and a Rossmann fold that interacts with a dinucleotide positioned for re-hydride transfer (Suppl. Fig. 4, A). In the OCD of *P. putida*, oligomerization results in a 14-stranded, closed β-barrel and each subunit contributes residues as shown in the Suppl. Fig. 4,A) and Tab. 1. In the PipA_*Sp*_ structure the β-barrel is similarly structured (Suppl. Fig. 4, B, and Suppl. Fig. 5, A; Tab. 1).

**Tab 1.**
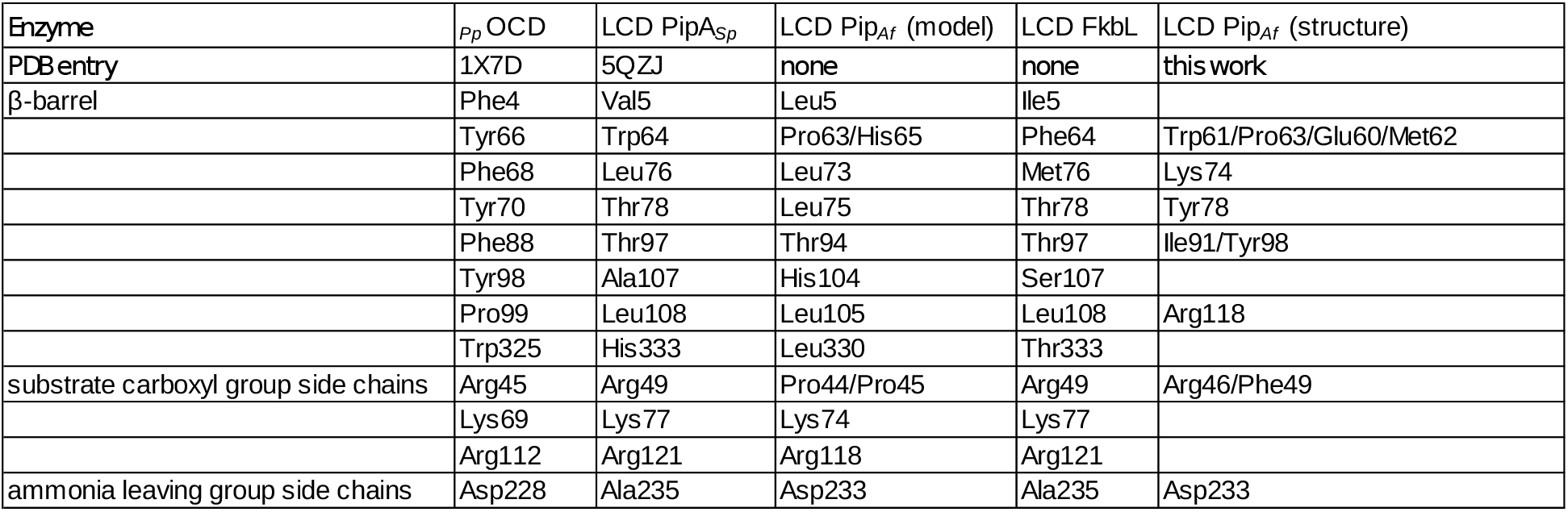
Comparision of key amino acid residues in the catalytic domain of the ornithine cyclodeaminase _*Pp*_OCD (1×7D) (Goodman et al., 2004), and the lysine cyclodeaminase PipA_*Sp*_ (5QZJ) (Ying, 2016), as well as the models of Pip_*Af*_, model of FkbL and the crystal structure of Pip_*Af*_ (this work).

Based on the crystal structures of _*Pp*_OCD and PipA_*Sp*_, the model structures of other LCDs Pip_*Af*_ and FkbL were generated through homology modelling. The structures of PipA_*Sp*_, Pip_*Af*_ and FkbL were overlaid; and the identified amino acid residues of PipA_*Sp*_ were used to find the corresponding key amino acids in the catalytic center of Pip_*Af*_ and FkbL. Comparisons with PipA_*Sp*_ demonstrated that in the lysine cyclodeaminase Pip_*Af*_ from *A. friuliensis* (Fig. 5) the β-barrel is similarly but not identically structured (Suppl. Fig. 5, B, C; Tab. 1). In the lysine cyclodeaminase FkbL from *S. tsukubaensis* the composition β-barrel differs from Pip_*Af*_, but is almost identical to PipA_*Sp*_ (Suppl. Fig. 6, A, B; Tab. 1).

**Fig. 5.**
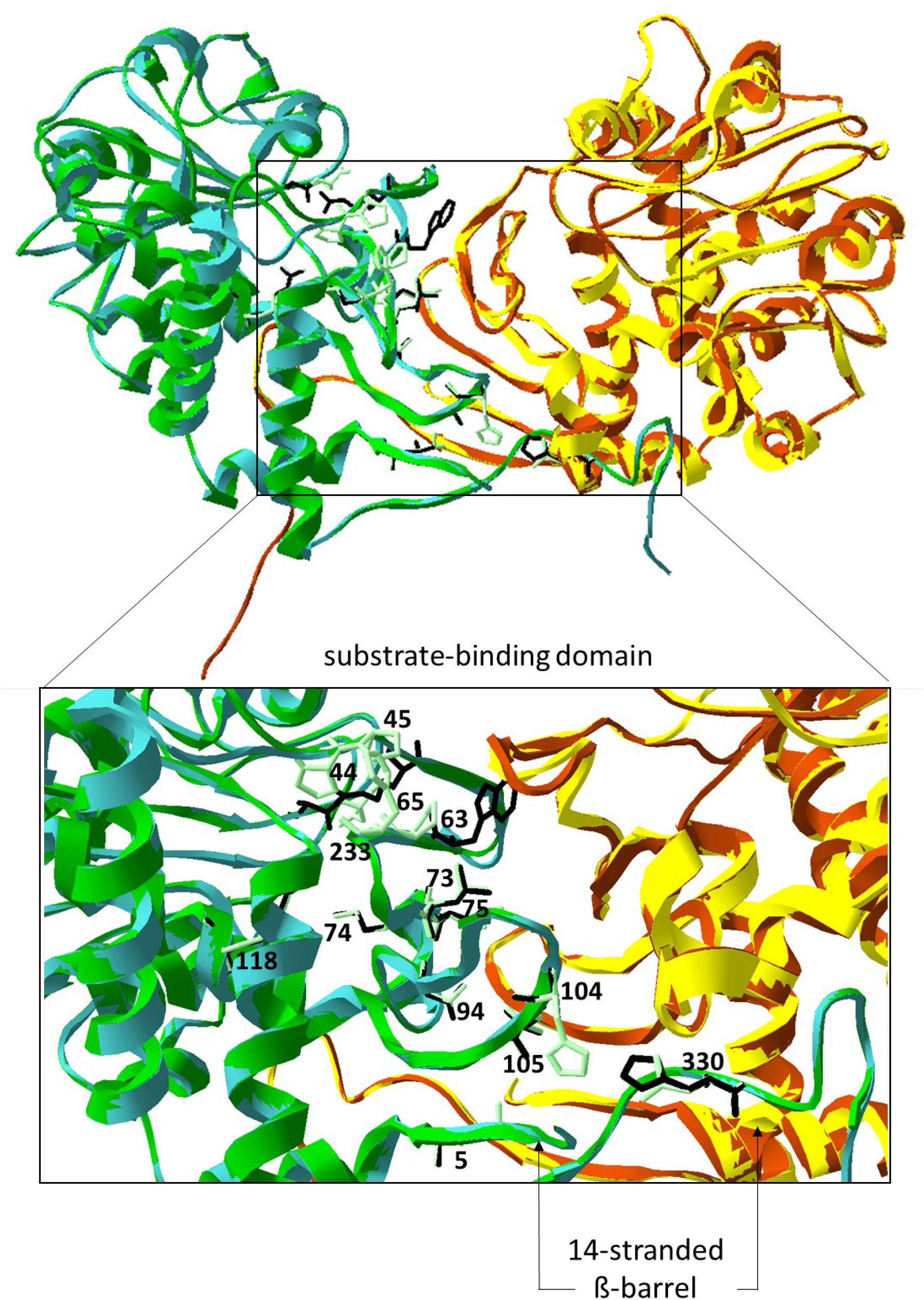
Model structure of Pip_*Af*_. The dimer is depicted in green (first monomer) and yellow (second monomer)) superposed with the crystal structure of the lysine cyclodeaminase PipA_*Sp*_ (the dimer in cyan (first monomer) and brown (second monomer)). Amino acid residues of Pip_*Af*_ marked in light green and labeled, of PipA_*Sp*_ in black and non-labeled. Amino acid residues are demonstrated on one monomer. Zoom-in represents the substrate binding domain with the β-barrel (7-stranded on each monomer).

Our studies of the crystal structure of the lysine cyclodeaminase PipA_*Sp*_ from *S. pristinaespiralis* as well as model structures of FkbL from *S. tsukubaensis* and Pip_*Af*_ from *A. friuliensis* revealed structural differences between Pip_*Af*_ and PipA_*St*_/FkbL. Key amino acids in Pip_*Af*_ and FkbL models were identified showing that Pip_*Af*_ features different amino acid residues in the catalytic domain (Fig. 5; Tab. 1).

### Determination of the crystal structure of Pip_*Af*_ from *A. friuliensis* and its structural analysis

Homology modelling demonstrated that the composition of Pip_*Af*_ differs from PipA_*Sp*_ and FkbL. In order to allow the engineering and specific design of Pip_*Af*_ for optimized FK-506 production, we aimed to solve its crystal structure. Pip_*Af*_ was overexpressed as a His-tagged protein in *E. coli* BL21 and purified for further analysis. A monomer of Pip_*Af*_ consists of 339 residues and the purified enzyme appeared as a ∼36 kDa protein in SDS-page. Its oligomeric state from size exclusion chromatography was found to be dimeric as observed for many members of the µ-crystallin/OCD family, e.g. _*Pp*_OCD (Goodman et al., 2004). Pip_*Af*_ crystals grew within few hours and diffracted up to 1.3 Å resulting in excellent crystallographic statistics (Suppl. Table 1). The enzyme consists of a homodimeric fold (Fig. 6) whose subunits include two domains that function in the L-lysine and NAD+/H cofactor binding (substrate binding domain) as well as in oligomerisation of subunits. One molecule NADH binds per monomer via a canonical Rossman fold, consistent with previous reports for the structurally characterized _*Pp*_OCD (Goodman et al., 2004; Dzurov et al., 2015).

**Fig. 6.**
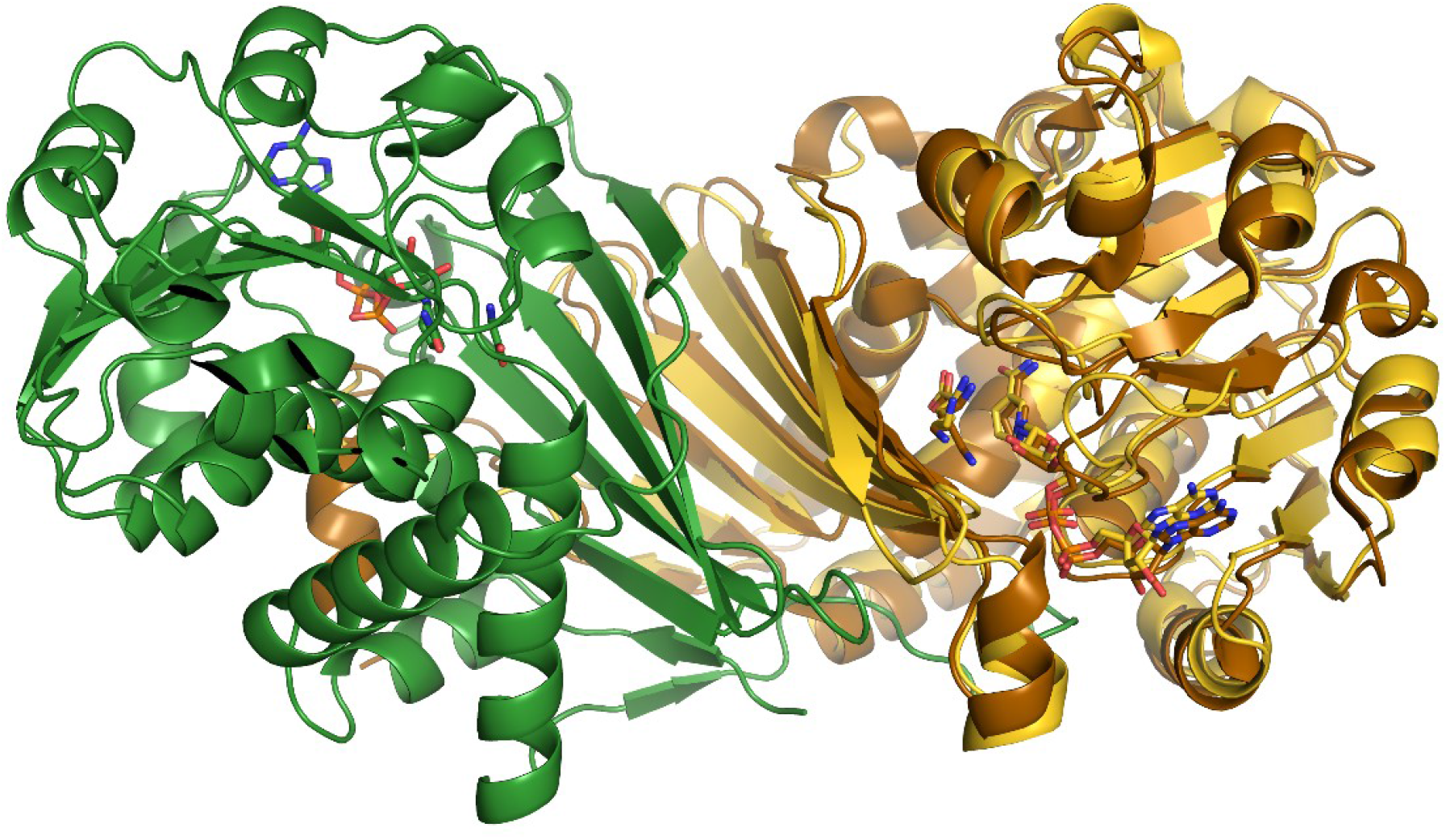
Pip_*Af*_ with NADH and lysine (dimer in yellow and green) superposed with OCD with NADH and ornithine (brown)

A 7-stranded β-sheet domain forms on the one side an extended dimer interface of the Pip_*Af*_ LCD, on the other side it contributes most of the residues of the substrate binding pocket (Fig. 6). The large interface between two monomer subunits with a surface of 2996 Å2 confers the stability of the dimer with -45 kcal/mol as calculated by the PISA server. Interestingly, both subunits contribute alternating hydrophobic residues (Phe49, Trp61, Pro63 and Tyr98) to the dimer interface and lock it similar to zipper-folds.

The model of Pip_*Af*_ was compared with the obtained crystal structure of Pip_*Af*_. The analysis of amino acid residues of interest in the catalytic site of the enzyme revealed that residues Pro63 and Asp233 could be confirmed via crystallographic studies to be involved into the key functions of the catalytic domain of the enzyme.

### Analysis of lysine binding pockets in the substrate binding domain of Pip_*Af*_

In order to enhance the properties of Pip_*Af*_ we were aiming to engineer the enzyme by site-directed mutagenesis. In order to identify relevant amino acids involved in substrate binding, _*Pp*_OCD crystal structure was compared with the obtained Pip_*Af*_ crystal structure. The substrate binding domain contributes conserved residues essential for conversion of L-lysine to L-pipecolate and directs them closely to NAD_+_. Like in _*Pp*_OCD, series of the acidic and basic residues interacting with lysine are highly conserved in Pip_*Af*_ (Arg46, Glu60, Met62, Lys74, Tyr78, Ile91, Arg118, Asp233). The domain is composed of a varied α / β fold involving a 7-stranded β-sheet packed against three α helices. Lysine binds in the active site running parallel to the nicotinamide ring. Tyrosine 78 (Tyr78) and isoleucine 91 (Ile91) in the Pip_*Af*_ structure substitute smaller, but also hydrophobic residues glycine and valine in _*Pp*_OCD respectively. Such residues should allow the lysine binding (Fig.6, 7).

**Fig. 7.**
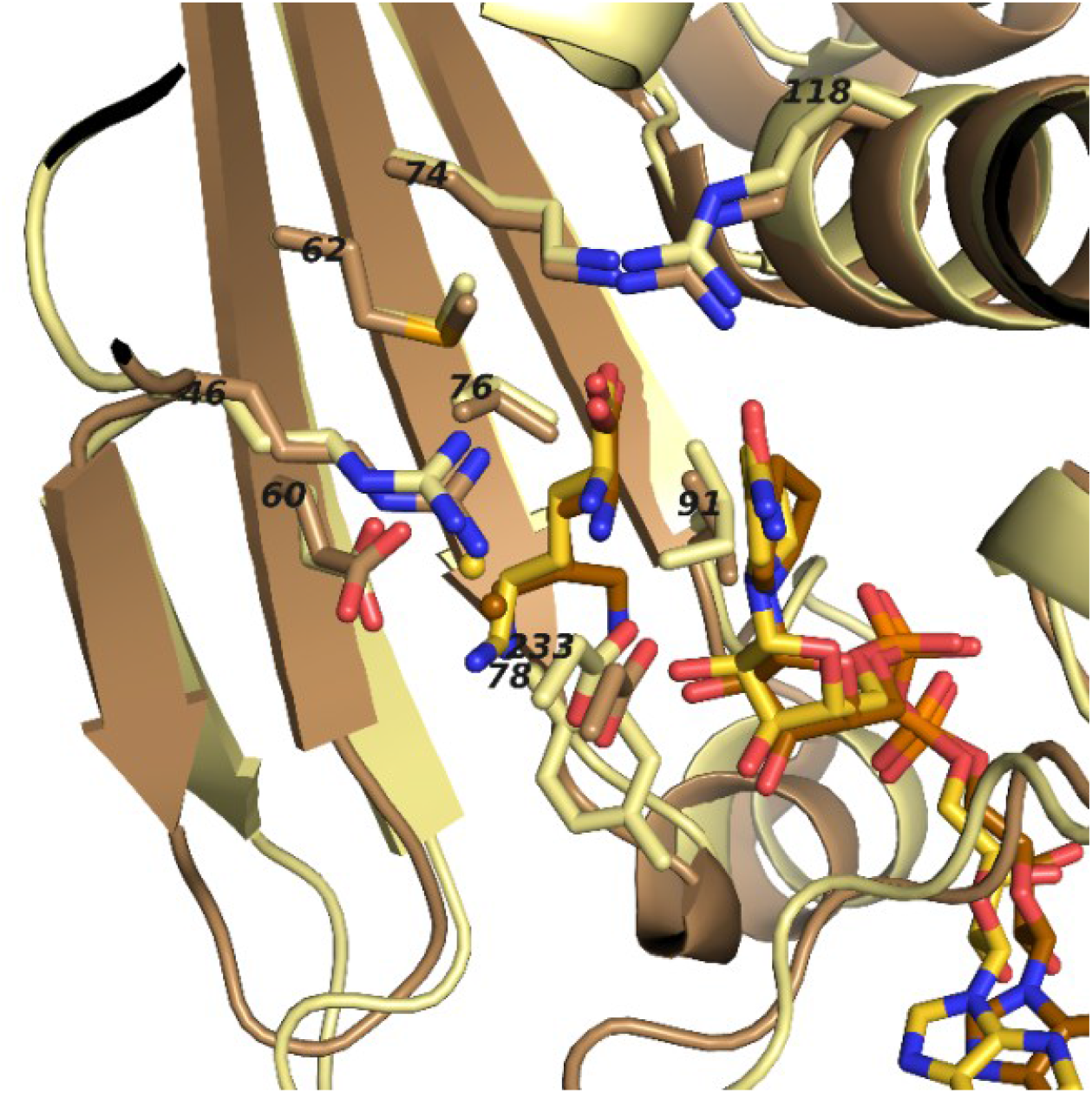
Superposed ligand binding pockets of Pip_*Af*_ with NADH and lysine (yellow) and _*Pp*_OCD with NADH and ornithine (brown). Residue numbers refer to Pip_*Af*_.

### Analysis of dinucleotide binding pocket in the substrate binding domain of Pip_*Af*_

Our attempts of Pip_*Af*_ engineering require in-depth understanding of the enzyme-dinucleotide binding kinetics. The catalytic mechanism of Pip_*Af*_ includes the dinucleotide cofactor binding at first, followed by substrate binding in an ordered reaction mechanism (Suppl. Fig. 7). All reported dinucleotide folds, also in _*Pp*_OCD, bind the pyrophosphoryl moiety of the dinucleotide via a glycine-rich, 30 to 35 amino-acids long loop comprising of a hydrophobic core, a negatively charged residue at the C-terminus and a positively charged residue at the N-terminus (Kleiger &Eisenberg, 2002; Goodman et al., 2004). A glycine-rich phosphate-binding sequence GxGxxG/A/S (where x is any residue) was found in all members of the µ-cristalline family, including _*Pp*_OCD (Goodman et al., 2004). Whereas most phosphate-binding motifs have the sequence GxGxxG/A, OCDs exhibit the sequence GxGxxS (Goodman et al., 2004). This explains why previous studies dismissed the existence of canonical dinucleotide binding sequences in OCDs (Kim et al., 1992). A precedent for use of the GxGxxS sequence was established in studies of dihydropyrimidine dehydrogenase, which uses the latter motif in FAD+ binding (Dobritzsch et al., 2001). The C-terminal part of Pip_*Af*_ lacks the coil-α6 segment reported to contribute some residues to NAD_+_/H recognition in OCD and thereby distinguish LCD from the rest of the μ-crystallin family (Goodman et al., 2004).

After purification by affinity chromatography and size exclusion chromatography Pip_*Af*_ exhibited an absorption at 340 nm corresponding to at least one molecule of NADH per dimer, if calculated with the extinction coefficient for NADH in aqueous solution. Crystallization was always performed with 1 mM NAD^+^, but each Pip_*Af*_ seems to contain NADH as cofactor as indicated by the non-planarity nicotinamide ring, which was clearly visible due to the high resolution. In analogy to the proposed mechanism of OCD the catalytic cycle of Pip_*Af*_ should start with a NAD^+^ oxidizing the alpha-amino group of lysine. The resulting imine group is substituted by the epsilon-amino group of lysine and the circular imine is finally reduced to pipicolic acid, thereby regenerating the NAD^+^ cofactor. OCD also contained NADH, therefore we would like to propose an alternative 2-step mechanism for cyclodeamination in which a direct substitution of the alpha-amino group by the hydride ion from NADH takes place which in turn is substituted by the epsilon-amino group.

### Site directed mutagenesis of key amino acids of Pip _*Af*_ to enhance the production of pipecolic acid

In order to optimize the one-step LCD-mediated synthesis of pipecolic acid through metabolic engineering we aimed to carry out site-directed mutagenesis based on our structural studies. The mutations included the exchange of amino acid residues in the substrate binding domain, potentially leading to enhanced substrate binding capacities, including binding pocket residues: Val58 by Leu58, Val58 by Ala58, Glu60 by Ala60, Glu60 by Gln60, Glu60 by Leu60 (Fig. 8, red; Tab. 1). The replacement of Val58 by Leu58 or Ala58 would lead to the strengthening or attenuation of the formation of a bond to the lysine side chain, which could be beneficial for deprotonation. The replacement Glu60 by Ala60 or Leu60 might open the entrance to the binding pocket a little more for the substrate. The replacement Glu60 by Gln60 might result in the generation of another hydrogen bond. Another strategy was the exchange of amino acid residues in the ammonia leaving group side chain residue Asp233 by Asn233 (Fig. 8, red; Tab. 1). Such substitution might lead to the generation of another hydrogen bond as well. Especially interesting should be the exchange of the lysine binding residue Ile91 by Val91. Ile at this place corresponds to the Val in FkbL. IIle binds hydrophobically the middle part of lysine, unlike Val. In order to avoid the reduction in activity due to this fact, Ile91 was replaced through Val. (Fig. 8, light green).

**Fig. 8.**
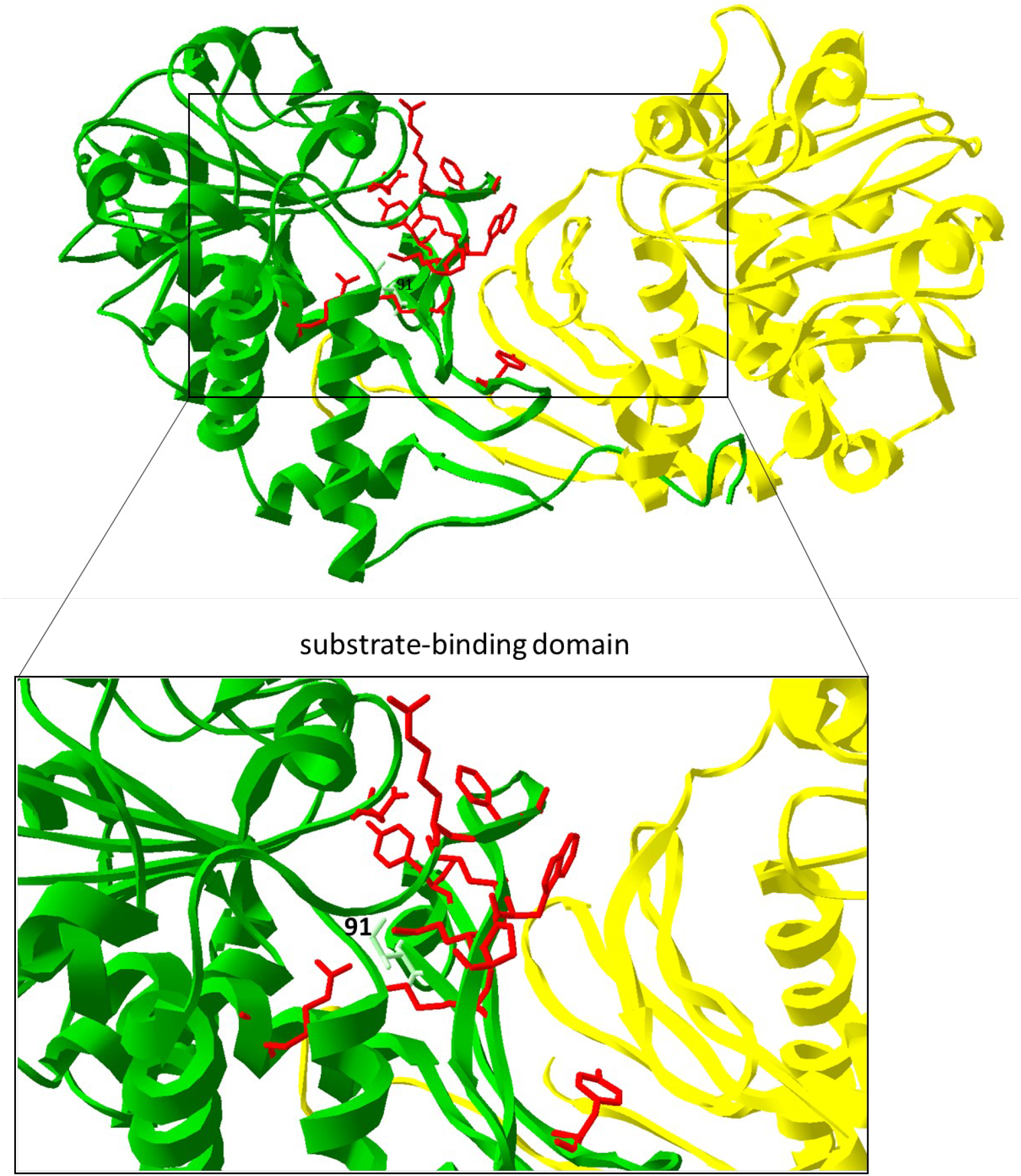
Engineering of Pip_*Af*_. Key amino acid residues in the substrate-binding domain of Pip_*Af*_ are marked in red; the replacement of Ile91 by Val is marked in light green and labeled. The dimer is shown in green (first monomer) and yellow (second monomer). Amino acid residues are demonstrated on one dimer. Zoom-in represents the substrate binding domain (7-stranded β-barrel on each monomer).

The selected amino acids were exchanged by site-specific mutagenesis in the pET30/*pip* expression plasmid. Using specifically designed primers with mutated bases, the mutagenesis PCR was carried out. The degradation of the non-mutated template DNA was performed with the restriction enzyme D*pn*I. Afterwards, the competent *E. coli* Novablue strain was transformed. Plasmid-carrying clones were identified by selection for kanamycin and verifyed by sequencing.

Mutated versions of the *pip*_*Af*_ gene (*pip*\*Ile91Val, *pip*\*Val58Leu, *pip*\*Val58Ala) were cloned into the expression vector and introduced into *E. coli* Rosetta 2 (DE3) pLysS. The product of the lysine cyclodeaminase reaction was detected via thin layer chromatography (TLC). In order to investigate and compare the activity of the individual mutants of the Pip_*Af*_ enzyme, a qualitative detection system for pipecolate production was established. The native *pip* gene from *A. friuliensis* served as reference. Since the host *E. coli* Rosetta 2 (DE3) pLysS has no lysine cyclodeaminase, the strain is not able to synthesize pipecolic acid and was therefore used as a negative control.

TLC revealed that lysine could only be converted into pipecolic acid by the *E*.*coli* Rosetta strains carrying the plasmids pET30/*pip*\*Ile91Val, pET30/*pip*\*Val58Leu, pET30/*pip*\*Val58Ala and the native pET30/*pip* with the native *pip*_*Af*_ gene (Fig. 9). In the mutated Pip variants Pip*Glu60Ala, Pip*Glu60Gln and Pip*Glu60Leu only a minimal pipecolic acid production was detected. In the case of the Pip* variant Pip*Asp233Asn, no pipecolic acid formation was observed. In the case of the Pip* variants Pip*Ile91Val and Pip*Val58Leu, the highest pipecolic acid formation was determined (Fig. 9).

**Fig. 9.**
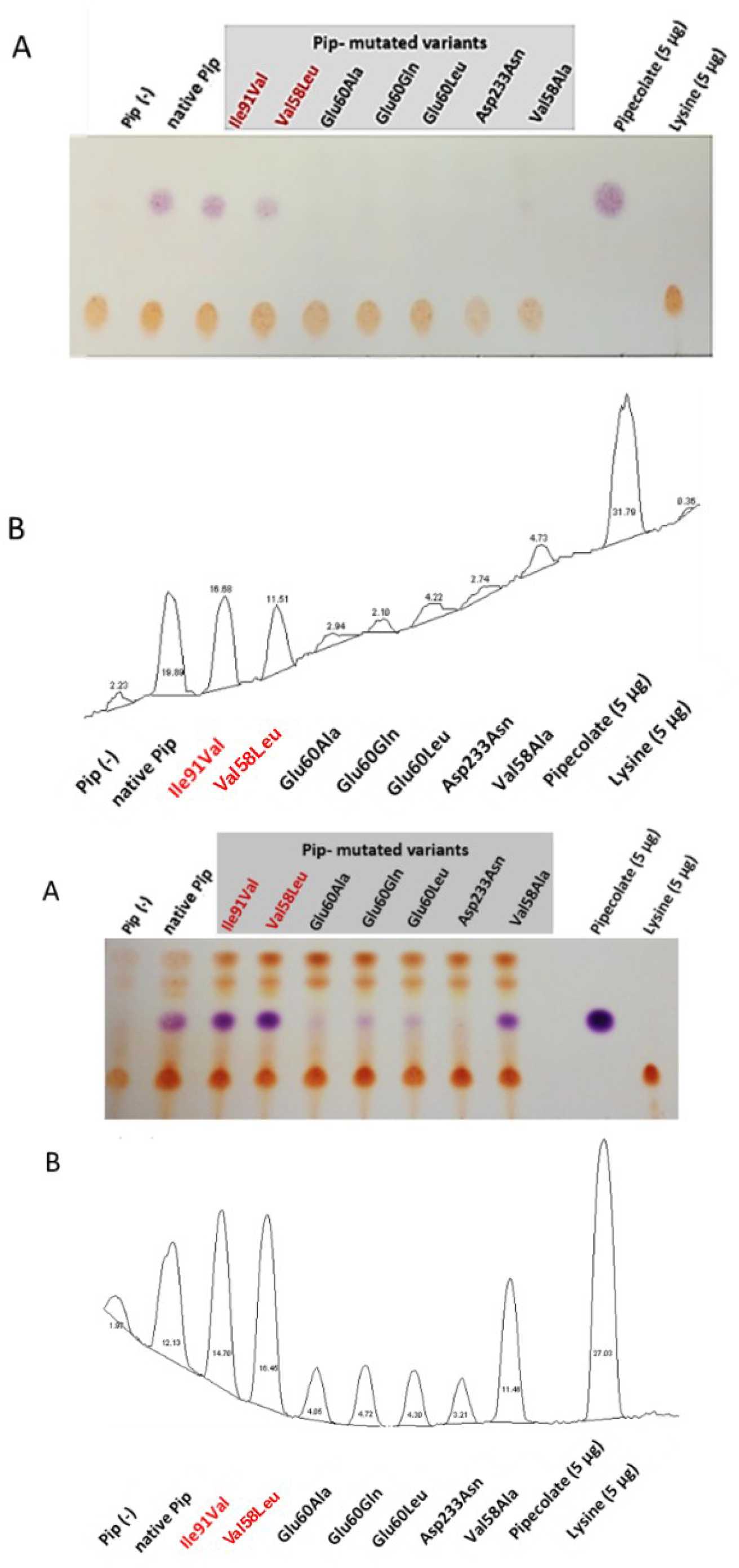
Proof of the pipecolate production in recombinant *E. coli* from two independent experiments. Sample injection: 5 µL of the culture supernatant. A: Scan of the TLC plate (contrast optimized). Violet drops represent pipecolate, yellow dots: lysine. Mutants with the best pipecolate production are marked in red: Ile91Val and Val58Leu.. B: Relative quantification of pipecolate and lysine based on digital imaging (ImageJ). Percentage of each peak is related to the total area of all peaks. Each peak corresponds to the respective spot on the TLC plate.

### FK-506 production by metabolically engineered strains

According to the crystal structure of Pip_*Af*_, Val91 does not hydrophobically bind the middle part of lysine, which should lead to enhanced enzyme activity. Thus, the mutated variant of Pip*Ile91Val was selected for the introduction into the recombinant host. The plasmid carrying the mutation pET30/*pip*\*Ile91Val was generated by site-directed mutagenesis through the introduction of the point mutation using specific primers. *S. tsukubaensis* with the previously introduced *dapA*_*St*_ and *lysC*_*Cg*_ genes was used as host. The specific FK-506 production was quantified using RP-HPLC and represented in mg/g cells and in percentage. While the *S. tsukubaensis* WT could achieve a FK-506 production of 1.7 mg/g cells, the *S. tsukubaensis* strain with the introduced mutated variant of Pip_*Af*_ produced of 3.4 mg/g, which is about 2-fold higher compared to the WT strain (Fig. 10). Again, lysine supplementation did not result in an FK-506 yield increase in the engineered strain *S. tsukubaensis* pET30/*pip*\*Ile91Val, which correlates with studies showing that the availability of nutrients like lysine can have an inhibitory effect on antibiotic production (van Wezel and McDowall, 2011).

**Fig. 10.**
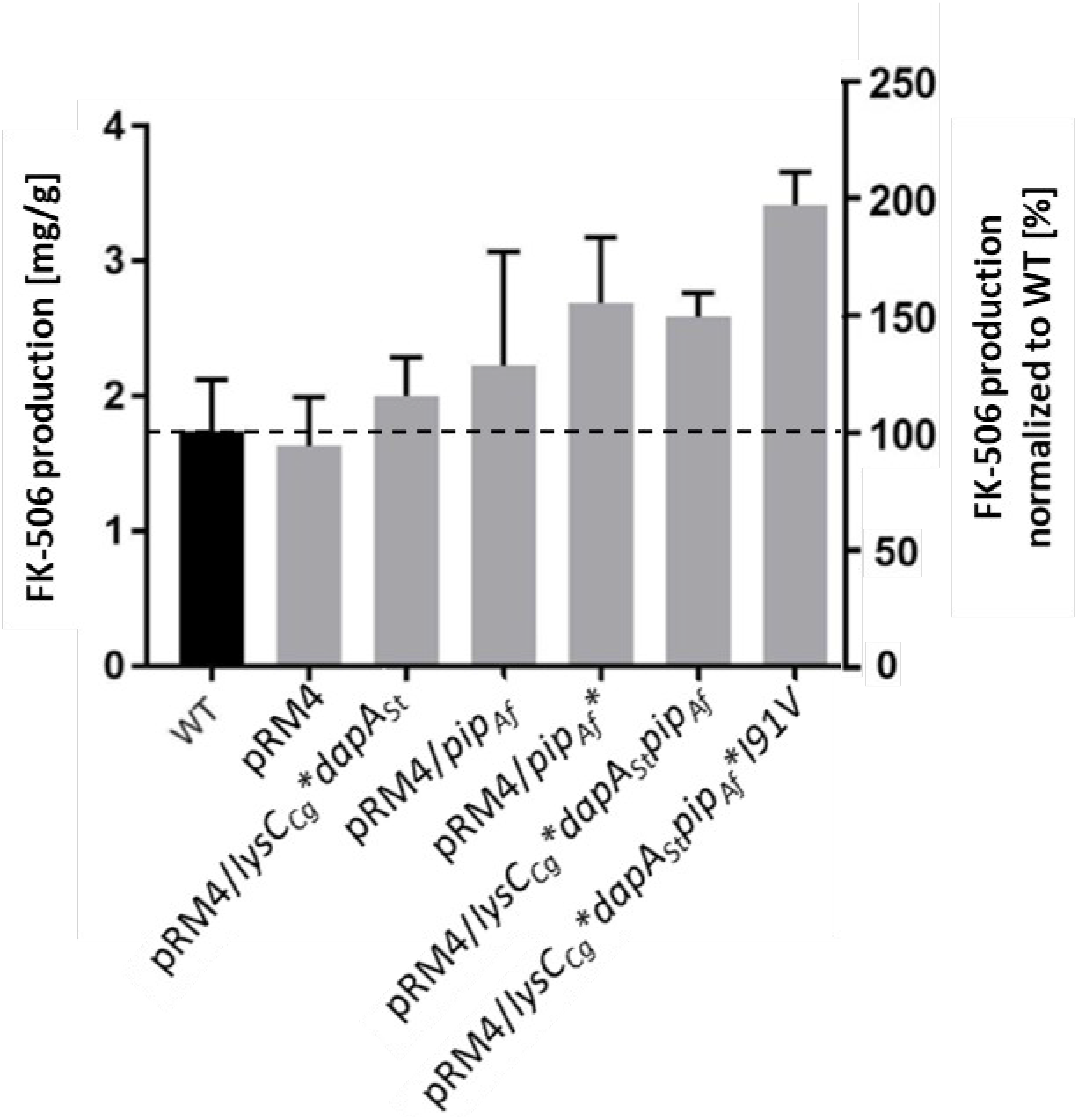
Production of FK-506 in the *S. tsukubaensis* as well as in engineered strains. X-axis on right: percentage representation of production optimization; as 100% the FK-506 production of the wild type (WT) was set at the respective time. X-axis on left: calculation of the mean of specific tacrolimus production over a fermentation period of six days. pRM4 stays for the integrative plasmid without an inserted gene. pRM4/ - integration of an additional copy of the gene on the integrative plasmide pRM4 encoding: *dapA*_*St*_ - dihydropicolinate synthase; *lysC*_*Cg*_*\** – deregulated aspartate kinase; *pip*_*Af*_ – LCD from *A. friuliensis*; *pip*_*Af*_*\*I91V -* LCD from *A. friuliensis* with Ile91 exchanged to Val. A T-test showed a p-value of 0.0007.

## Conclusions

The complex biosynthesis of FK-506 in the wild type producer *S. tsukubaensis* only results in rather low titers. However, this biosynthesis includes key steps that can be open to attack. We could show that engineering two steps of FK-506 production are suitable targets for optimization: the provision of lysine which is required as precursor for the FK-506 building block pipecolate, and the conversion of lysine into pipecolate.

In order to increase the lysine pool, engineering of two reactions of the primary metabolism turned out to be successful: on the one hand directing the metabolic flux from aspartate to lysine was possible by overexpression of the dihydropicolinate synthase gene. On the other hand a synthetic biology approach using the gene for a feedback deregulated aspartate kinase from a lysine overproducing *C. glutamicum* resulted in a yield improvement.

The lysine converting lysine cyclodeaminase (LCD) was shown to be essential for generating pipecolate. Overexpression of the *S. tsukubaensis* LCD gene as well as that of LCD genes from other actinomycetes, which synthetize pipecolate-containing natural products, delivered recombinant *S. tsukubaensis* strains with increased FK-506 yields. Since the different LCDs provided different levels of optimization, we concluded that metabolic engineering of LCD may offer an additional option for further increase. Indeed, after crystallization of the LCD from *A. friuliensis* it was possible to identify amino acids, the exchange of which by site-directed mutagenesis resulted in improved FK-506 production.

Whereas the individual engineering steps led to moderate yield increase, the combination of overexpression of biosynthetic genes with enzyme design and with a synthetic biology approach resulted in a duplication of the FK-506 yield.

## Authorship contribution statement

S.S. performed studies of the *S. tsukubaensis lysC*-*dapA* strain. C.S. and T.S. solved the crystal structure of Pip_*Af*_. S.K. performed homology modelling of Pip_*Af*_, PipA_*Sp*_, FkbL, comparisons of amino acid sequences and protein structures. A.B. and W.W. designed the study. A.B. and S.K. contributed to writing of the manuscript. S.S., A.B., S.K., C.S. and C.B. prepared the figures. C.B. performed TLC analysis as well as FK-506 quantification and tests with the final engineered strain. W.W. edited the manuscript and provided helpful feedback.

## Acknowledgements

We would like to thank Andreas Kulik for the support with HPLC analysis of the FK-506. This work was funded by the Bundesministerium für Bildung und Forschung (BMBF) (FKZ 031B0269A), in the frame of the ERA-CoBioTech project “Tacrodrugs”. We are grateful to our partners from the ERA-CoBioTech consortium for helpful discussions. We would also like to acknowledge the support by the BMBF Fördermaßnahmen “Targetvalidierung für die pharmazeutische Wirkstoffentwicklung” (project GPS-TBT (FKZ: 16 GW0183K) as well as project GSS-TUBTAR (FKZ: 16 GW0253K)) and the Deutsche Forschungsgemeinschaft (DFG, German Research Foundation) project ID398967434 (TRR 261, TPs B01 and B08).

## Literature

1. Altschul, S. F., Madden, T. L., Schäffer, A. A., Zhang, J., Zhang, Z., Miller, W., & Lipman, D. J. 1997. Gapped BLAST and PSI-BLAST: a new generation of protein database search programs. Nucleic Acids Res 25, 3389–3402. https://doi.org/10.1093/nar/25.17.3389

2. Andexer, J. N., Kendrew, S. G., Nur-e-Alam, M., Lazos, O., Foster, T. A., Zimmermann, A. S., Warneck, T. D., Suthar, D., Coates, N. J., Koehn, F. E., Skotnicki, J. S., Carter, G. T., Gregory, M. A., Martin, C. J., Moss, S. J., Leadlay, P. F., & Wilkinson, B. 2011. Biosynthesis of the immunosuppressants FK506, FK520, and rapamycin involves a previously undescribed family of enzymes acting on chorismate. Proc Natl Acad Sci U S A 108, 4776–4781. https://doi.org/10.1073/pnas.1015773108

3. Aretz, W., Meiwes, J., Seibert, G., Vobis, G., & Wink, J. 2000. Friulimicins: novel lipopeptide antibiotics with peptidoglycan synthesis inhibiting activity from Actinoplanes friuliensis sp. nov. 4Taxonomic studies of the producing microorganism and fermentation. J Antibiot (Tokyo) 53, 807–815. https://doi.org/10.7164/antibiotics.53.807

4. Ban, Y. H., Park, S. R., & Yoon, Y. J. 2016. The biosynthetic pathway of FK506 and its engineering: from past achievements to future prospects. J Ind Microbiol Biotechnol 43, 389–400. https://doi.org/10.1007/s10295-015-1677-7

5. Baranasic, D., Gacesa, R., Starcevic, A., Zucko, J., Blazic, M., Horvat, M., Gjuracic, K., Fujs, S., Hranueli, D., Kosec, G., Cullum, J., & Petkovic, H. 2013. Draft Genome Sequence of Streptomyces rapamycinicus Strain NRRL 5491, the Producer of the Immunosuppressant Rapamycin. Genome Announc. 1 e00581–13. https://doi.org/10.1128/genomeA.00581-13

6. Barreiro, C., & Martínez-Castro, M. 2014. Trends in the biosynthesis and production of the immunosuppressant tacrolimus (FK506). Appl Microbiol Biotechnol 98, 497–507. https://doi.org/10.1007/s00253-013-5362-3

7. Barreiro, C., Prieto, C., Sola-Landa, A., Solera, E., Martínez-Castro, M., Pérez-Redondo, R., García-Estrada, C., Aparicio, J. F., Fernández-Martínez, L. T., Santos-Aberturas, J., Salehi-Najafabadi, Z., Rodríguez-García, A., Tauch, A., & Martín, J. F. 2012. Draft genome of Streptomyces tsukubaensis NRRL 18488, the producer of the clinically important immunosuppressant tacrolimus (FK506). J Bacteriol 194, 3756–3757. https://doi.org/10.1128/JB.00692-12

8. Bis, D. M., Ban, Y. H., James, E. D., Alqahtani, N., Viswanathan, R., & Lane, A. L. 2015. Characterization of the nocardiopsin biosynthetic gene cluster reveals similarities to and differences from the rapamycin and FK-506 pathways. Chembiochem 16, 990–997. https://doi.org/10.1002/cbic.201500007

9. Blazic, M., Starcevic, A., Lisfi, M., Baranasic, D., Goranovic, D., Fujs, S., Kuscer, E., Kosec, G., Petkovic, H., Cullum, J., Hranueli, D., & Zucko, J. 2012. Annotation of the modular polyketide synthase and nonribosomal peptide synthetase gene clusters in the genome of Streptomyces tsukubaensis NRRL18488. Appl Environ Microbiol 78, 8183–8190. https://doi.org/10.1128/AEM.01891-12

10. Borel, F., Hachi, I., Palencia, A., Gaillard, M. C., & Ferrer, J. L. 2014. Crystal structure of mouse mu-crystallin complexed with NADPH and the T3 thyroid hormone. FEBS J 281, 1598–1612. https://doi.org/10.1111/febs.12726

11. Byun, S.M., Jeong, S.W., Cho, D.H., Kim Y.H. 2015. Optimized conversion of L-lysine to L-pipecolic acid using recombinant lysine cyclodeaminase from Streptomyces pristinaespiralis. Biotechnol Bioproc E 20, 73–78. https://doi.org/10.1007/s12257-014-0428-3

12. Chen, D., Zhang, L., Pang, B., Chen, J., Xu, Z., Abe, I., & Liu, W. 2013. FK506 maturation involves a cytochrome p450 protein-catalyzed four-electron C-9 oxidation in parallel with a C-31 O-methylation. J Bacteriol 195, 1931–1939. https://doi.org/10.1128/JB.00033-13

13. Chen, D., Zhang, Q., Zhang, Q., Cen, P., Xu, Z., & Liu, W. 2012. Improvement of FK506 production in Streptomyces tsukubaensis by genetic enhancement of the supply of unusual polyketide extender units via utilization of two distinct site-specific recombination systems. Appl Environ Microbiol 78, 5093–5103. https://doi.org/10.1128/AEM.00450-12

14. Cheng, Y. R., Fang, A., & Demain, A. L. 1995. Effect of amino acids on rapamycin biosynthesis by Streptomyces hygroscopicus. Appl Microbiol Biotechnol 43, 1096–1098. https://doi.org/10.1007/BF00166931

15. Cocito C. 1979. Antibiotics of the virginiamycin family, inhibitors which contain synergistic components. Microbiol Rev 43, 145–192. https://doi.org/10.1128/mr.43.2.145-192.1979

16. Dobritzsch, D., Schneider, G., Schnackerz, K. D., & Lindqvist, Y. 2001. Crystal structure of dihydropyrimidine dehydrogenase, a major determinant of the pharmacokinetics of the anti-cancer drug 5-fluorouracil. EMBO J 20, 650–660. https://doi.org/10.1093/emboj/20.4.650

17. Dzurová, L., Forneris, F., Savino, S., Galuszka, P., Vrabka, J., & Frébort, I. 2015. The three-dimensional structure of “Lonely Guy” from Claviceps purpurea provides insights into the phosphoribohydrolase function of Rossmann fold-containing lysine decarboxylase-like proteins. Proteins 83, 1539–1546. https://doi.org/10.1002/prot.24835

18. Fu, L. F., Tao, Y., Jin, M. Y., & Jiang, H. 2016. Improvement of FK506 production by synthetic biology approaches. Biotechnol Lett 38, 2015–2021. https://doi.org/10.1007/s10529-016-2202-4

19. Gatto, G. J., Jr, Boyne, M. T., 2nd, Kelleher, N. L., & Walsh, C. T. 2006. Biosynthesis of pipecolic acid by RapL, a lysine cyclodeaminase encoded in the rapamycin gene cluster. J Am Chem Soc 128, 3838–3847. https://doi.org/10.1021/ja0587603

20. Goodman, J. L., Wang, S., Alam, S., Ruzicka, F. J., Frey, P. A., & Wedekind, J. E. 2004. Ornithine cyclodeaminase: structure, mechanism of action, and implications for the mu-crystallin family. Biochemistry 43, 13883–13891. https://doi.org/10.1021/bi048207i

21. Goranovic, D., Kosec, G., Mrak, P., Fujs, S., Horvat, J., Kuscer, E., Kopitar, G., & Petkovic, H. 2010. Origin of the allyl group in FK506 biosynthesis. J Biol Chem 285, 14292–14300. https://doi.org/10.1074/jbc.M109.059600

22. Guex, N., Peitsch, M. C., & Schwede, T. 2009. Automated comparative protein structure modeling with SWISS-MODEL and Swiss-PdbViewer: a historical perspective. Electrophoresis 30 Suppl 1, S162–S173. https://doi.org/10.1002/elps.200900140

23. Gupta, R. N., & Spenser, I. D. 1969. Biosynthesis of the piperidine nucleus. The mode of incorporation of lysine into pipecolic acid and into piperidine alkaloids. J Biol Chem 244, 88–94.

24. Haddad, E. M., McAlister, V. C., Renouf, E., Malthaner, R., Kjaer, M. S., & Gluud, L. L. 2006. Cyclosporin versus tacrolimus for liver transplanted patients. Cochrane Database Syst Rev (4), CD005161. https://doi.org/10.1002/14651858.CD005161.pub2

25. He M., 2006. Pipecolic acid in microbes: biosynthetic routes and enzymes. J Ind Microbiol Biotechnol 33, 401–407. https://doi.org/10.1007/s10295-006-0078-3

26. Huang D., Li S., Xia M., Wen J., Jia X. 2013. Genome-scale metabolic network guided engineering of Streptomyces tsukubaensis for FK506 production improvement. Microb Cell Fact 12:52.

27. Huang, D., Li, S., Xia, M., Wen, J., & Jia, X. 2013. Genome-scale metabolic network guided engineering of Streptomyces tsukubaensis for FK506 production improvement. Microb Cell Fact 12, 52. https://doi.org/10.1186/1475-2859-12-52

28. Jiang, H., & Kobayashi, M. 1999. Differences between cyclosporin A and tacrolimus in organ transplantation. Transplant Proc 31, 1978–1980. https://doi.org/10.1016/s0041-1345(99)00235-3

29. Jiang, H., Haltli, B., Feng, X., Cai, P., Summers, M., Lotvin, J., & He, M. 2011. Investigation of the biosynthesis of the pipecolate moiety of neuroprotective polyketide meridamycin. J Antibiot (Tokyo) 64, 533–538. https://doi.org/10.1038/ja.2011.45

30. Jung, S., Moon, S., Lee, K., Park, Y. J., Yoon, S., & Yoo, Y. J. 2009. Strain development of Streptomyces sp. for tacrolimus production using sequential adaptation. J Ind Microbiol Biotechnol 36, 1467–1471. https://doi.org/10.1007/s10295-009-0634-8

31. Kieser T., Bibb M.J., Buttner M.J., Chater K.F., Hopwood D.A. 2000. Practical Streptomyces genetics. John Innes Foundation, Norwich

32. Kim, R. Y., Gasser, R., & Wistow, G. J. 1992. mu-crystallin is a mammalian homologue of Agrobacterium ornithine cyclodeaminase and is expressed in human retina. Proc Natl Acad Sci U S A 89, 9292–9296. https://doi.org/10.1073/pnas.89.19.9292

33. Kim, H. S., & Park, Y. I. 2007. Lipase activity and tacrolimus production in Streptomyces clavuligerus CKD 1119 mutant strains. J Microbiol Biotechnol 17, 1638–1644.

34. Kim, H. S., & Park, Y. I. 2008. Isolation and identification of a novel microorganism producing the immunosuppressant tacrolimus. J Biosci Bioeng 105, 418–421. https://doi.org/10.1263/jbb.105.418

35. Kino, T., Hatanaka, H., Hashimoto, M., Nishiyama, M., Goto, T., Okuhara, M., Kohsaka, M., Aoki, H., & Imanaka, H. 1987. FK-506, a novel immunosuppressant isolated from a Streptomyces. I. Fermentation, isolation, and physico-chemical and biological characteristics. The Journal of antibiotics 40, 1249–1255. https://doi.org/10.7164/antibiotics.40.1249

36. Kleiger, G., & Eisenberg, D. 2002. GXXXG and GXXXA motifs stabilize FAD and NAD(P)-binding Rossmann folds through C(alpha)-H… O hydrogen bonds and van der waals interactions. J Antibiot (Tokyo) 323, 69–76. https://doi.org/10.1016/s0022-2836(02)00885-9

37. Müller, C., Nolden, S., Gebhardt, P., Heinzelmann, E., Lange, C., Puk, O., Welzel, K., Wohlleben, W., & Schwartz, D. 2007. Sequencing and analysis of the biosynthetic gene cluster of the lipopeptide antibiotic Friulimicin in Actinoplanes friuliensis. Antimicrob Agents Chemother 51, 1028–1037. https://doi.org/10.1128/AAC.00942-06

38. Li, Y., Liang, S., Wang, J., Ma, D., & Wen, J. 2019. Enhancing the production of tacrolimus by engineering target genes identified in important primary and secondary metabolic pathways and feeding exogenous precursors. Bioprocess Biosyst Eng 42, 1081–1098. https://doi.org/10.1007/s00449-019-02106-9

39. Lin A. N. 2010. Innovative use of topical calcineurin inhibitors. Dermatol Clin 28, 535–545. https://doi.org/10.1016/j.det.2010.03.008

40. Liu, J., Farmer, J. D., Jr, Lane, W. S., Friedman, J., Weissman, I., & Schreiber, S. L. 1991. Calcineurin is a common target of cyclophilin-cyclosporin A and FKBP-FK506 complexes. Cell 66, 807–815. https://doi.org/10.1016/0092-8674(91)90124-h

41. Martínez-Castro, M., Salehi-Najafabadi, Z., Romero, F., Pérez-Sanchiz, R., Fernández-Chimeno, R. I., Martín, J. F., & Barreiro, C. 2013. Taxonomy and chemically semi-defined media for the analysis of the tacrolimus producer ‘Streptomyces tsukubaensis’. Appl Microbiol Biotechnol 97, 2139–2152. https://doi.org/10.1007/s00253-012-4364-x

42. Mast, Y., Weber, T., Gölz, M., Ort-Winklbauer, R., Gondran, A., Wohlleben, W., & Schinko, E. 2011. Characterization of the ‘pristinamycin supercluster’ of Streptomyces pristinaespiralis. Microb Biotechnol 4, 192–206. https://doi.org/10.1111/j.1751-7915.2010.00213.x

43. Mendelovitz, S., & Aharonowitz, Y. 1983. beta-lactam antibiotic production by Streptomyces clavuligerus mutants impaired in regulation of aspartokinase. J Gen Microbiol 129, 2063–2069. https://doi.org/10.1099/00221287-129-7-2063

44. Mendelovitz, S., & Aharonowitz, Y. 1982. Regulation of cephamycin C synthesis, aspartokinase, dihydrodipicolinic acid synthetase, and homoserine dehydrogenase by aspartic acid family amino acids in Streptomyces clavuligerus. Antimicrob Agents Chemother 21, 74–84. https://doi.org/10.1128/AAC.21.1.74

45. Menges, R., Muth, G., Wohlleben, W., & Stegmann, E. 2007. The ABC transporter Tba of Amycolatopsis balhimycina is required for efficient export of the glycopeptide antibiotic balhimycin. Appl Microbiol Biotechnol 77, 125–134. https://doi.org/10.1007/s00253-007-1139-x

46. He M. 2006. Pipecolic acid in microbes: biosynthetic routes and enzymes. Journal of industrial microbiology & biotechnology 33, 401–407. https://doi.org/10.1007/s10295-006-0078-3

47. Mo, S., Kim, D. H., Lee, J. H., Park, J. W., Basnet, D. B., Ban, Y. H., Yoo, Y. J., Chen, S. W., Park, S. R., Choi, E. A., Kim, E., Jin, Y. Y., Lee, S. K., Park, J. Y., Liu, Y., Lee, M. O., Lee, K. S., Kim, S. J., Kim, D., Park, B. C., … Yoon, Y. J. 2011. Biosynthesis of the allylmalonyl-CoA extender unit for the FK506 polyketide synthase proceeds through a dedicated polyketide synthase and facilitates the mutasynthesis of analogues. J Ind Microbiol Biotechnol 133, 976–985. https://doi.org/10.1021/ja108399b

48. Mohammadpour, N., Elyasi, S., Vahdati, N., Mohammadpour, A. H., & Shamsara, J. 2011. A review on therapeutic drug monitoring of immunosuppressant drugs. Iran J Basic Med Sci 14, 485–498.

49. Motamedi, H., Cai, S. J., Shafiee, A., & Elliston, K. O. 1997. Structural organization of a multifunctional polyketide synthase involved in the biosynthesis of the macrolide immunosuppressant FK506. Eur J Biochem 244, 74–80. https://doi.org/10.1111/j.1432-1033.1997.00074.x

50. Motamedi, H., & Shafiee, A. 1998. The biosynthetic gene cluster for the macrolactone ring of the immunosuppressant FK506. Eur J Biochem 256, 528–534. https://doi.org/10.1046/j.1432-1327.1998.2560528.x

51. Motamedi, H., Shafiee, A., Cai, S. J., Streicher, S. L., Arison, B. H., & Miller, R. R. 1996. Characterization of methyltransferase and hydroxylase genes involved in the biosynthesis of the immunosuppressants FK506 and FK520. J Bacteriol 178, 5243–5248. https://doi.org/10.1128/jb.178.17.5243-5248.1996

52. Muduma, G., Saunders, R., Odeyemi, I., & Pollock, R. F. 2016. Systematic Review and Meta-Analysis of Tacrolimus versus Ciclosporin as Primary Immunosuppression After Liver Transplant. PLoS One 11, e0160421. https://doi.org/10.1371/journal.pone.0160421

53. Ordóñez-Robles, M., Santos-Beneit, F., & Martín, J. F. 2018. Unraveling Nutritional Regulation of Tacrolimus Biosynthesis in Streptomyces tsukubaensis through omic Approaches. Antibiotics (Basel) 7, 39. https://doi.org/10.3390/antibiotics7020039

54. Özcengiz, G., Okay, S., Ünsaldı, E., Taşkın, B., Liras, P., & Piret, J. 2010. Homologous expression of aspartokinase (ask) gene in Streptomyces clavuligerus and its hom-deleted mutant: effects on cephamycin C production. Bioeng Bugs 1, 191–197. https://doi.org/10.4161/bbug.1.3.11244

55. Park, S. R., Yoo, Y. J., Ban, Y. H., & Yoon, Y. J. 2010. Biosynthesis of rapamycin and its regulation: past achievements and recent progress. J Antibiot (Tokyo) 63, 434–441. https://doi.org/10.1038/ja.2010.71

56. Pérez-García, F., Peters-Wendisch, P., & Wendisch, V. F. 2016. Engineering Corynebacterium glutamicum for fast production of L-lysine and L-pipecolic acid. Appl Microbiol Biotechnol 100, 8075–8090. https://doi.org/10.1007/s00253-016-7682-6

57. Ramachandran G.N., Ramakrishnan C., Sasisekharan V. 1963. Stereochemistry of polypeptide chain configurations. J Mol Biol 7, 95–99. https://doi.org/10.1016/s0022-2836(63)80023-6

58. Qian, Z. G., Xia, X. X., & Lee, S. Y. 2011. Metabolic engineering of Escherichia coli for the production of cadaverine: a five carbon diamine. Biotechnol Bioeng 108, 93–103. https://doi.org/10.1002/bit.22918

59. Sahm, H., Eggeling, L., Eikmanns, B., & Krämer, R. 1996. Construction of L-lysine-, L-threonine-, and L-isoleucine-overproducing strains of Corynebacterium glutamicum. Ann N Y Acad Sci 782, 25–39. https://doi.org/10.1111/j.1749-6632.1996.tb40544.x

60. Sambrook J., Russell R. Molecular cloning: a laboratory manual 2001. 3rd ed. Cold Spring Harbor Laboratory Press, New York, NY.

61. Schreiber, S. L., & Crabtree, G. R. 1992. The mechanism of action of cyclosporin A and FK506. Immunol Today 13, 136–142. https://doi.org/10.1016/0167-5699(92)90111-J

62. Seibold, G., Auchter, M., Berens, S., Kalinowski, J., & Eikmanns, B. J. 2006. Utilization of soluble starch by a recombinant Corynebacterium glutamicum strain: growth and lysine production. J Biotechnol 124, 381–391. https://doi.org/10.1016/j.jbiotec.2005.12.027

63. Staatz, C. E., & Tett, S. E. 2004. Clinical pharmacokinetics and pharmacodynamics of tacrolimus in solid organ transplantation. Clin Pharmacokinet 43, 623–653. https://doi.org/10.2165/00003088-200443100-00001

64. Steinmetz, H., Glaser, N., Herdtweck, E., Sasse, F., Reichenbach, H., & Höfle, G. 2004. Isolation, crystal and solution structure determination, and biosynthesis of tubulysins--powerful inhibitors of tubulin polymerization from myxobacteria. Angew Chem Int Ed Engl 43, 4888–4892. https://doi.org/10.1002/anie.200460147

65. Storz, G., & Imlay, J. A. 1999. Oxidative stress. Curr Opin Microbiol 2, 188–194. https://doi.org/10.1016/s1369-5274(99)80033-2

66. Studer, G., Rempfer, C., Waterhouse, A. M., Gumienny, R., Haas, J., & Schwede, T. 2020. QMEANDisCo-distance constraints applied on model quality estimation. Bioinformatics 36, 1765–1771. https://doi.org/10.1093/bioinformatics/btz828

67. Taber, D. J., Dupuis, R. E., Fann, A. L., Andreoni, K. A., Gerber, D. A., Fair, J. H., Johnson, M. W., & Shrestha, R. 2002. Tacrolimus dosing requirements and concentrations in adult living donor liver transplant recipients. Liver Transpl 8, 219–223. https://doi.org/10.1053/jlts.2002.30885

68. Tani, Y., Miyake, R., Yukami, R., Dekishima, Y., China, H., Saito, S., Kawabata, H., & Mihara, H. 2015. Functional expression of L-lysine α-oxidase from Scomber japonicus in Escherichia coli for one-pot synthesis of L-pipecolic acid from DL-lysine. Appl Microbiol Biotechnol 99, 5045–5054. https://doi.org/10.1007/s00253-014-6308-0

69. Temuujin, U., Chi, W. J., Lee, S. Y., Chang, Y. K., & Hong, S. K. 2011. Overexpression and biochemical characterization of DagA from Streptomyces coelicolor A3(2): an endo-type β-agarase producing neoagarotetraose and neoagarohexaose. Appl Microbiol Biotechnol 92(4), 749–759. https://doi.org/10.1007/s00253-011-3347-7

70. Tsotsou, G. E., & Barbirato, F. 2007. Biochemical characterisation of recombinant Streptomyces pristinaespiralis L-lysine cyclodeaminase. Biochimie 89, 591–604. https://doi.org/10.1016/j.biochi.2006.12.008

71. Turło, J., Gajzlerska, W., Klimaszewska, M., Król, M., Dawidowski, M., & Gutkowska, B. 2012. Enhancement of tacrolimus productivity in Streptomyces tsukubaensis by the use of novel precursors for biosynthesis. Enzyme Microb Technol 51, 388–395. https://doi.org/10.1016/j.enzmictec.2012.08.008

72. van Wezel, G. P., & McDowall, K. J. 2011. The regulation of the secondary metabolism of Streptomyces: new links and experimental advances. Natural product reports 28(7), 1311–1333. https://doi.org/10.1039/c1np00003a

73. Watanabe, S., Tozawa, Y., & Watanabe, Y. 2014. Ornithine cyclodeaminase/μ-crystallin homolog from the hyperthermophilic archaeon Thermococcus litoralis functions as a novel Δ(1)-pyrroline-2-carboxylate reductase involved in putative trans-3-hydroxy-l-proline metabolism. FEBS Open Bio 4, 617–626. https://doi.org/10.1016/j.fob.2014.07.005

74. Waterhouse, A., Bertoni, M., Bienert, S., Studer, G., Tauriello, G., Gumienny, R., Heer, F. T., de Beer, T., Rempfer, C., Bordoli, L., Lepore, R., & Schwede, T. 2018. SWISS-MODEL: homology modelling of protein structures and complexes. Nucleic Acids Res 46, W296–W303. https://doi.org/10.1093/nar/gky427

75. Wilkinson, C. J., Hughes-Thomas, Z. A., Martin, C. J., Böhm, I., Mironenko, T., Deacon, M., Wheatcroft, M., Wirtz, G., Staunton, J., & Leadlay, P. F. 2002. Increasing the efficiency of heterologous promoters in actinomycetes. J Mol Microbiol Biotechnol 4, 417–426.

76. Wu, K., Chung, L., Revill, W. P., Katz, L., & Reeves, C. D. 2000. The FK520 gene cluster of Streptomyces hygroscopicus var. ascomyceticus (ATCC 14891) contains genes for biosynthesis of unusual polyketide extender units. Gene 251, 81–90. https://doi.org/10.1016/s0378-1119(00)00171-2

77. Xia, M., Huang, D., Li, S., Wen, J., Jia, X., & Chen, Y. 2013. Enhanced FK506 production in Streptomyces tsukubaensis by rational feeding strategies based on comparative metabolic profiling analysis. Biotechnol Bioeng 110, 2717–2730. https://doi.org/10.1002/bit.24941

78. Ying H.X., Wang J., Wang Z., Feng J., Chen K.Q., Li Y., Ouyang P. 2015. Enhanced conversion of L-lysine to L-pipecolic acid using a recombinant Escherichia coli containing lysine cyclodeaminase as whole-cell biocatalyst. J Mol Catal B-Enzym 117, 75–80

79. Ying, H., Tao, S., Wang, J., Ma, W., Chen, K., Wang, X., & Ouyang, P. 2017. Expanding metabolic pathway for de novo biosynthesis of the chiral pharmaceutical intermediate L-pipecolic acid in Escherichia coli. Microb Cell Fact 16, 52. https://doi.org/10.1186/s12934-017-0666-0

80. Ying, H.; Chen, K. 2016. 5GZJ PDB data base entry Cyclodeaminase_PA

81. Yoon, Y., & Choi, C. 1997. Nutrient effects on FK-506, a new immunosuppressant, production by Streptomyces sp. in a defined medium. J Ferment Bioeng 83, 599–603.

82. Zheng, L., Baumann, U., & Reymond, J. L. 2004. An efficient one-step site-directed and site-saturation mutagenesis protocol. Nucleic Acids Res 32, e115. https://doi.org/10.1093/nar/gnh110

