## Supplementary figures for "Optimization of the precursor supply for an enhanced FK-506 production in *Streptomyces tsukubaensis*"

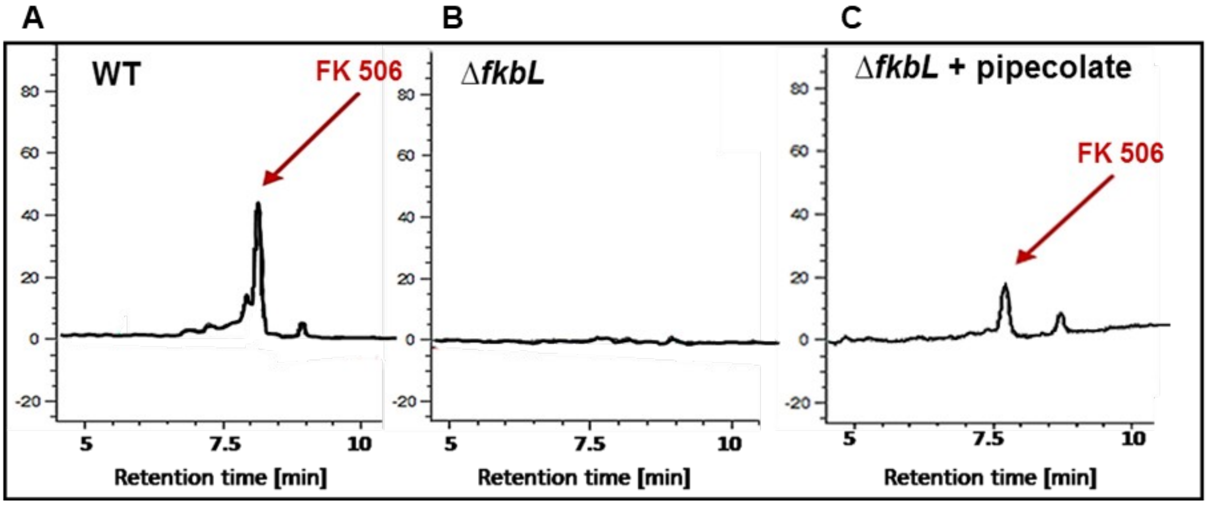


Supplementary Fig. 1. HPLC chromatograms for the detection of tacrolimus after three days of cultivation in MG medium. A) FK-506 in the *S. tsukubaensis* WT. B) no FK-506 in the *S. tsukubaensis ΔfkbL* mutant. C) FK-506 production in *S. tsukubaensis ΔfkbL* after the addition of pipecolate. X-axis: absorbance in mAU, y-axis: retention time (minutes).

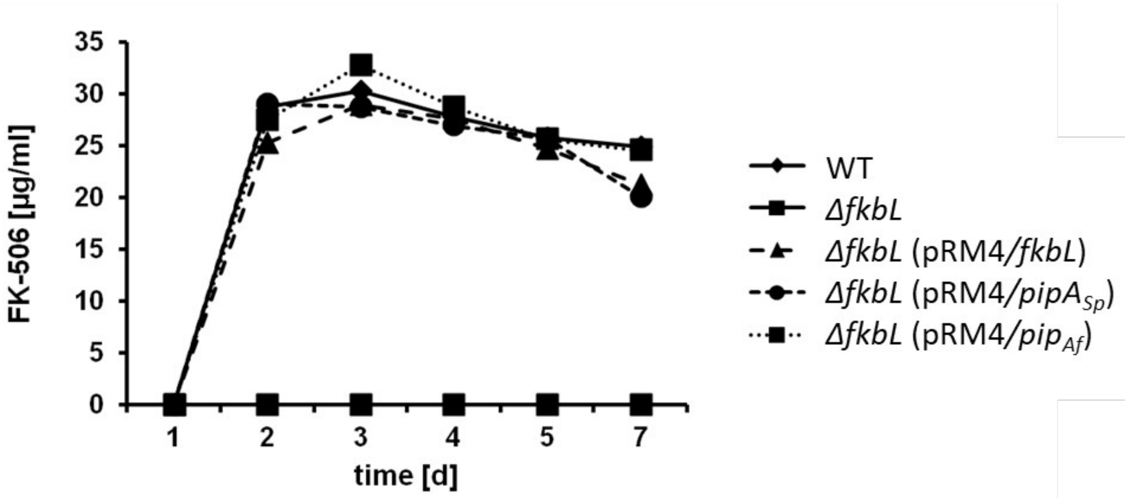


Supplementary Fig. 2. Genetic complementation of Δ*fkbL S.tsukubaensis* with lysine cylodeaminase genes from *S. tsukubaensis* WT, *S. pristinaespiralis* and *A. friuliensis*

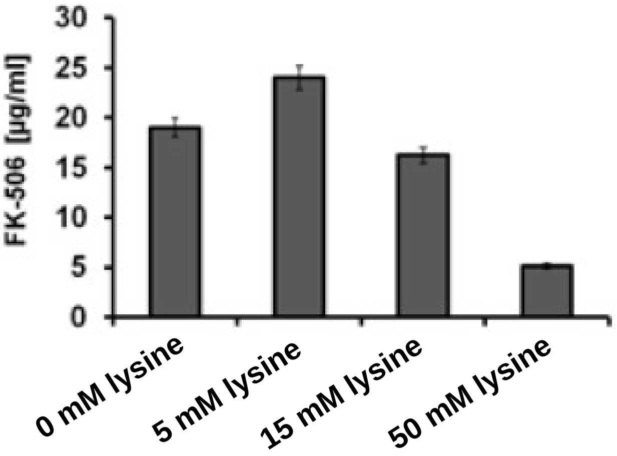


Supplementary Fig. 3. Production test of *S. tsukubaensis* WT in production medium MG with addition of different lysine concentrations. The exogenous supplementation of higher lysine concentrations leads to inhibited FK-506 production.

A

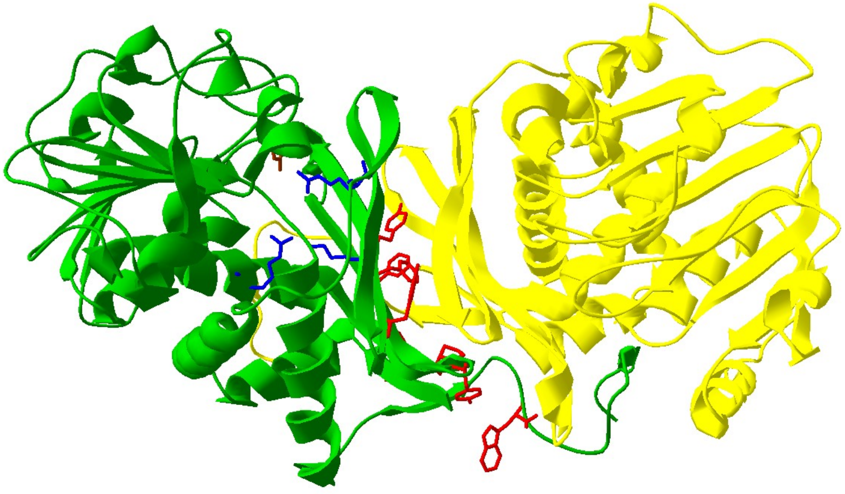

B

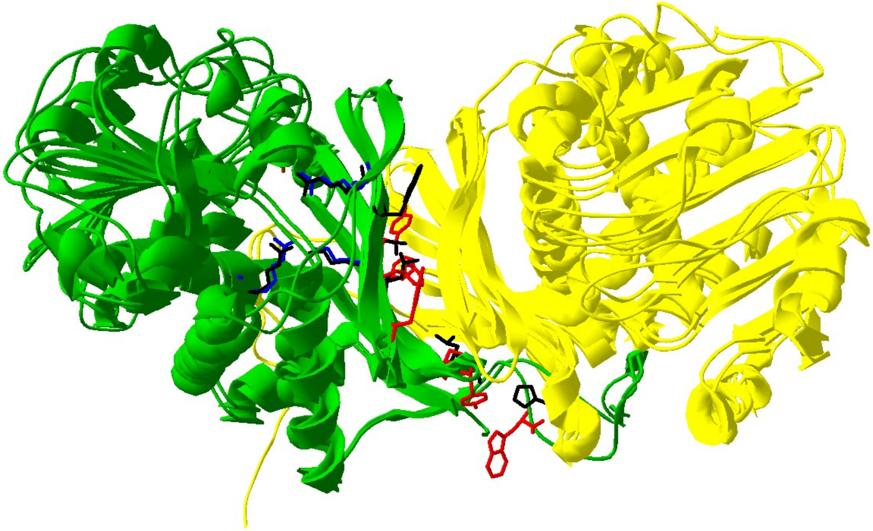

Suppl. Fig. 4. A. Structure of the ornithine cyclodeaminase from *Presudomonas putida* (1X7D), B. Comparision of the ornithine cyclodeaminase (1X7D) with the lysine cyclodeaminase from *Streptomyces pristinaespiralis* (5QZJ). Key amino acid residues are marked in red, blue and brown (1X7D) and black (5QZJ).

In the OCD of *P. putida* (1X7D), oligomerization results in a 14-stranded, closed β-barrel and each subunit contributes residues, namely Phe4, Tyr66, Phe68, Tyr70, Phe88, Tyr98, Pro99, and Trp325 (A, red) to the barrel interior in the substrate-binding domain. Moreover, the substrate carboxyl group interacts with the side chains of Arg45, Lys69, and Arg112 (A, blue) and the ammonia leaving group hydrogen bonds to the side chain of Asp228 (A, brown) (after Kim & Park, 2007; Goodman at al., 2004).

A

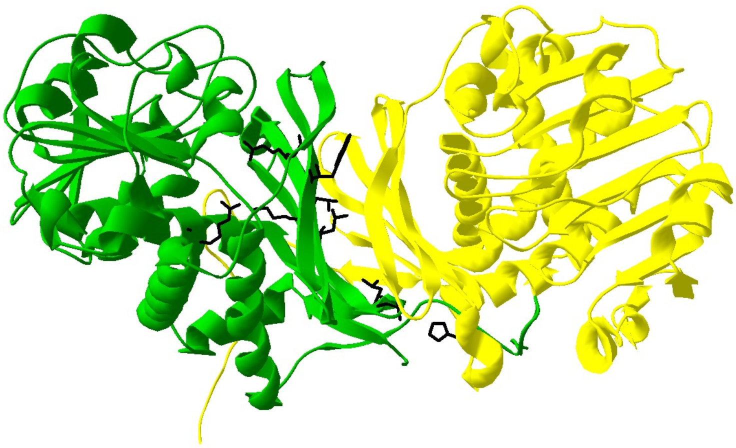

B

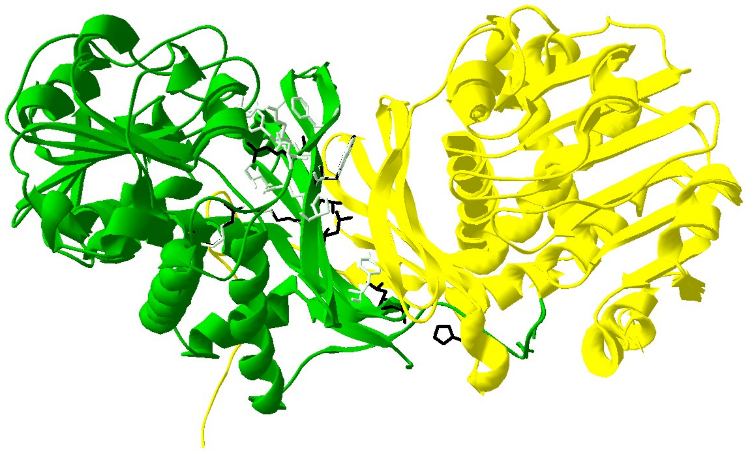

C

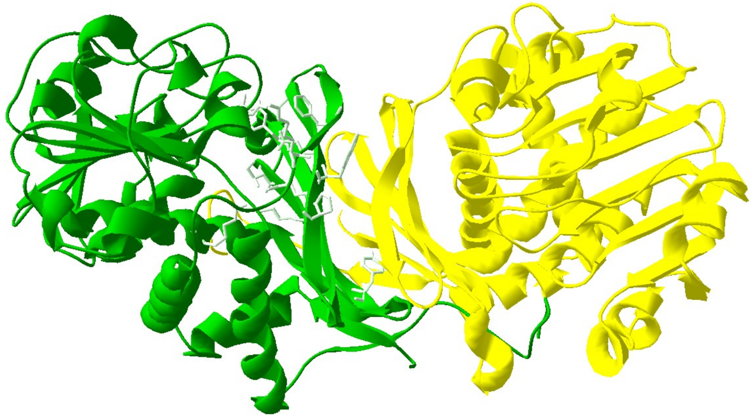

Suppl. Fig. 5. A. Structure of the lysine cyclodeaminase from *Streptomyces pristinaespiralis* (5QZJ); B. Comparision of the the lysine cyclodeaminase (5QZJ) with the model of the lysine cyclodeaminase Pip$_{Af}$, C. Model of the lysine cyclodeaminase from *Actinoplanes friuliensis.* Key amino acid residues are marked in black (5QZJ) and green (Pip$_{Af}$).

In the PipA$_{Sp}$ structure (5QZJ) the β-barrel is similarly structured involving at the same place residues Val5, Trp64, Leu76, Thr78, Thr97, Ala107, Leu108 and His333 compared to 1X7D. The substrate carboxyl group side chains include Arg49, Lys77 and Arg121 and the side chain for the ammonia leaving group the residue Ala235 (Suppl. Fig. 2, B, black; Suppl. Fig. 3, A). Comparisons with PipA$_{Sp}$ demonstrated that in the lysine cyclodeaminase Pip$_{Af}$ from *A. friuliensis* (Fig. 4) the β-barrel is similarly but not identically structured involving at the same place residues Leu5, Pro63/His65, Leu73, Leu75, Thr94, His104, Leu105, Leu330 (Suppl. Fig. 3, B, C, light green) compared to 5QZJ (Suppl. Fig. 3, A, B, black). The substrate carboxyl group side chains include Pro44/Pro45, Lys74, Arg118 and the side chain for the ammonia leaving group the residue Asp233 in the Pip$_{Af}$ structure (Suppl. Fig. 3, B, C, light green).

A

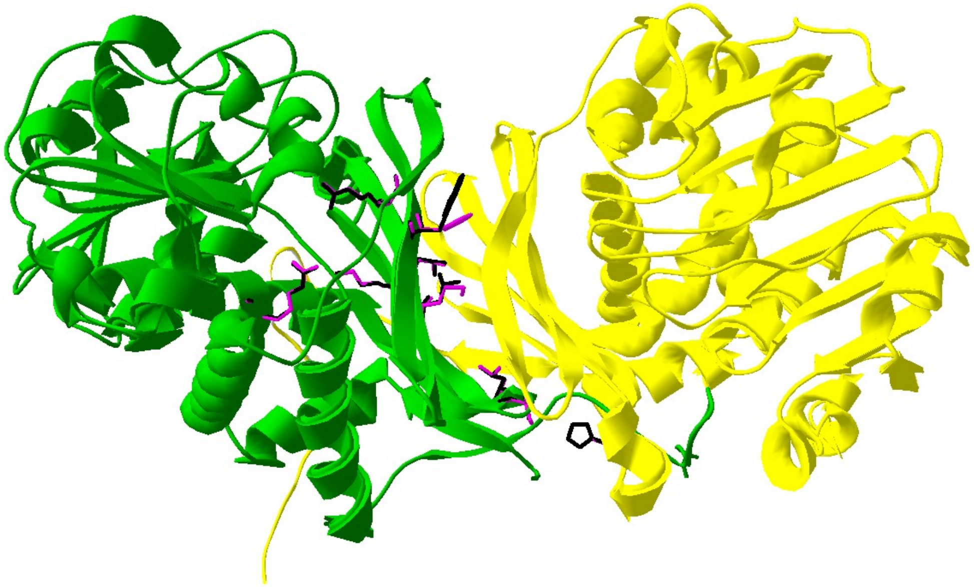

B

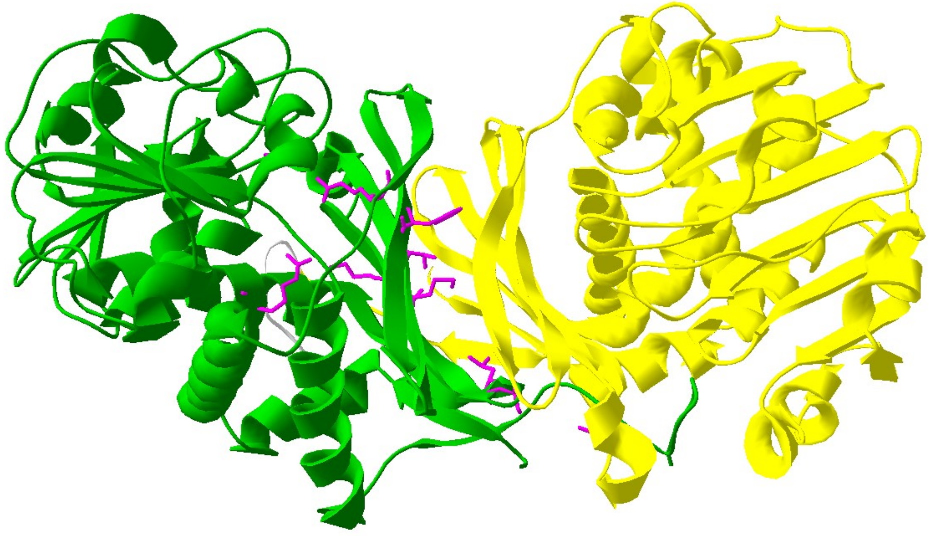

Suppl. Fig. 6. A. Comparision of the the lysine cyclodeaminase from *S. pristinaespiralis* (5QZJ) with the model of the lysine cyclodeaminase FkbL, B. Model of the lysine cyclodeaminase from *Streptomyces tsukubaensis.* Key amino acid residues are marked in black (5QZJ) and purple (FkbL).

In the lysine cyclodeaminase FkbL from *S. tsukubaensis* the composition β-barrel differs from $Pip_{Af}$, but is almost identical to $PipA_{Sp}$ involving at the same place residues Ile5, Phe64, Met76, Thr78, Thr97, Ser107, Leu108, Thr333 (Suppl. Fig. 4, A, B, purple) compared to 5QZJ (Suppl. Fig. 4, A, black). The substrate carboxyl group side chains include Arg49, Lys77, Arg121 and the side chain for the ammonia leaving group the residue Ala235 in the FkbL structure (Suppl. Fig. 4, A, B).

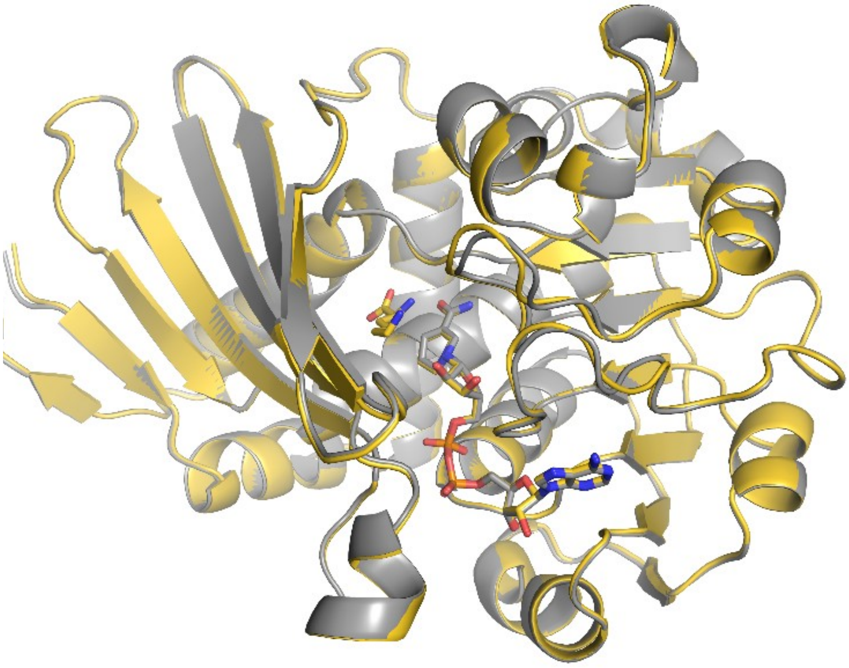

Suppl. Fig.7. $Pip_{Af}$ (yellow) with NADH and lysine superposed with $Pip_{Af}$ with only NADH (grey).

### Data collection statistics

| | **$Pip_{Af}$** | **$Pip_{Af}$ complex with Lys** |
|---|---|---|
| Beamline | SLS X06DA | SLS X06DA |
| Wavelength λ [Å] | 1.000 | 1.000 |
| Detector | Pilatus 2M | Pilatus 2M |
| Detector distance [mm] | 135 | 120 |
| Space group | $P2_12_12_1$ | $P2_12_12_1$ |
| Unit cell: [Å] | 59.2 89.9 137.3 | 59.5 89.2 139.1 |
| [degree] | 90.0 90.0 90.0 | 90.0 90.0 90.0 |
| Resolution [Å] | 50 – 1.4 (1.49 – 1.4) | 50.0 – 1.3 (1.38 – 1.3) |
| No. of reflections | | |
| Measured | 1318869 (195149) | 1306889 (186423) |
| Unique | 145265 (23022) | 181827 (28151) |
| $R_{meas}$ [%] | 10.3 (153.4) | 6.7 (112.7) |
| CC1/2 | 59.1 | 72.2 |
| Completeness (%) | 99.8 (98.8) | 99.3 (96.0) |
| Multiplicity | 9.1 (8.5) | 7.2 (6.6) |
| <I>/<σ(I)> | 12.1 (1.35) | 14.5 (1.62) |

| | **Pip$_{Af}$** | **Pip$_{Af}$ complex with Lys** |
|---|---|---|
| Wilson Factor [Å$^2$] | 24.2 | 22.9 |
| Crystal Mosaicity [°] | 0.076 | 0.083 |

**Refinement statistics**

| | **Pip$_{Af}$** | **Pip$_{Af}$ complex with Lys** |
|---|---|---|
| Resolution range [Å] | 50 – 1.3 | 50 – 1.4 |
| $R_{Cryst}$ | 0.219 | 0.205 |
| $R_{free}$ (test set of 5%) | 0.242 | 0.226 |
| No. of non-H atoms (partial occupancy) | | |
| chain A / chain B | 2552 / 2551 | 2552/2547 |
| NAD | 88 | 88 |
| lysine | - | 50 |
| water | 212 | 205 |
| Average isotropic B-Factor [Å$^2$] | 20.7 | 22.8 |
| main chain A / B | 24.7 / 18.7 | 20.6 / 18.4 |
| side chain A / B | 26.9 / 21.2 | 23.3 / 20.9 |
| NAD | 17.9 | 16.3 |
| lysine | - | 21.2 |
| water molecules | 23.5 | 23.0 |
| Rmsd for bond lengths [Å] | 0.006 | 0.006 |
| Rmsd for bond angle [°] | 1.15 | 1.11 |
| Ramachandran regions | | |
| most favorable [%] | 97.9 | 97.9 |
| allowed [%] | 1.9 | 2.0 |
| outliers [%] | 0.1 | 0 |

Suppl. Table. 1: Crystallographic data collection and refinement statistics

| Strains/Plasmids | Genotype | Reference |
|---|---|---|
| *E. coli* NovaBlue | recA1, endA1, gyrA96, thi-1, *hsdR17* (rK12 - ,mK12 + ) *supE44*, *relA1*, lac [F', *proAB*, *lacIq*, *lacZ*ΔM15, Tn10] (Tet$^{R}$) | Novagen |
| *E. coli* BL21(DE3)pLysS | F-, *ompT*, *hsdSB* (rB-mB-), *gal*, *dcm*, (DE3), pLysS, Cam R , methionine auxotroph, 35S-met labeling *lon*-, *ompT*-, *dcm*-, pLysS | Novagen |
| *E. coli* Rosetta 2(DE3) pLysS | Derivate of BL21, pRARE2: 7 rare tR-NAs; rare *E. coli* codons: arginine (AGA, AGG, CGA) glycine (GGA), isoleucine (AUA), leucine (CUA), pro-line (CCC) | Novagen |
| *E. coli* ET12567/pUZ8002 | Methylation deficient strain *E. coli* with pUZ8002, F-, dam- 13::Tn9, *dcm*-6, *hsdM*, *hsdR*, *lacY1*,Cam$^{R}$ ; Kan$^{R}$ | MacNeil et al.1992 |
| *E. coli* ET12567/pUB307 | Methylation deficient strain *E. coli* with pUB307, F-, dam-13::Tn9, *dcm*-6, *hsdM*, *hsdR*, *lacY1*, Cam$^{R}$ , Kan$^{R}$ , Tet$^{R}$ | Bennett et al., 1977; MacNeil et al., 1992 |
| *S. coelicolor* M145 | *S. coelicolor* A3(2) without native plasmids | Kieser et al., 2000 |
| *S. pristinaespiralis* Pr11 | Pristinamycin producer wild type | Aventis Pharma |
| *Streptomyces tsukubaensis* NRRL18488 | STP1 STP2 | Martinez-Castro et al., 2012 |
| *Streptomyces tsukubaensis* Δ*fkbL* | Derivate of *S. tsukubensis* WT; *fkbL* replaced by AprR; deficient in FK-506 production | This work |
| pRM4 | pSET152*ermEp** with artificial RBS, | Menges et al., 2007 |

| | | |
|---|---|---|
| | AprR | |
| pRM4/*fkbL* | pRM4 Derivate with *fkbL* (from *S. tsukubaensis*) | This work |
| pRM4/*fkbP* | pRM4 Derivate with *fkbP* (from *S. tsukubaensis*) | This work |
| pRM4/*fkbL fkbP* | pRM4 Derivate with *fkbL*, *ermE* and *fkbP* (from *S. tsukubaensis*) | This work |
| pRM4/*pipA* | pRM4 Derivate with *pipA* (from *S. pristinaespiralis*) | This work |
| pRM4/*pip* | pRM4 Derivate with *pip* (from *A. friuliensis*) | This work |
| pRM4/*pip* * E60A | pRM4 Derivate with *pip* * $^{E60A}$ | This work |
| pRM4/*pip* * I91V | pRM4 Derivate with *pip* * $^{I91V}$ | This work |
| pRM4/*pip* * D233N | pRM4 Derivate with *pip* * $^{D233N}$ | This work |
| pRM4/*pip* * L234A | pRM4 Derivate with *pip* * $^{L234A}$ | This work |
| pRM4 kan/*fkbL* | pRM4/*fkbL*-Derivate with additional $Kan^R$ | This work |
| pRM4 kan/*pipA* | pRM4/*pipA*-Derivate with additional $Kan^R$ | This work |
| pRM4 kan/*pip* | pRM4/*pip*-Derivate with additional $Kan^R$ | This work |
| pRM4/*ask* | pRM4-Derivate with *ask* (from *S. tsukubaensis*) | This work |
| pRM4/*ask* * | pRM4-Derivate with mutated *ask**S301Y (from *S. tsukubaensis*) | This work |
| pRM4/*lysC* * | pRM4-Derivate with deregulated *lysC** (from *C. glutamicum*) | This work |

| | | |
|---|---|---|
| pRM4/*dapA* | pRM4-Derivate with *dapA* (from *S. tsukubaensis*) | This work |
| pRM4/*dapA lysC* * | pRM4-Derivate with *dapA*, *ermE* und *lysC* * | This work |
| pSET152 | pUC18 *lacZα*, *oriT* (RK2), RP4 *mob* region, ΦC31 *int* and attP, $Apr^R$ | Bierman, et al., 1992; Schmitt-John & Engels,1992 |
| pDRIVE | T7 RNA-Polymerase Promotor, SP6 RNA-Polymerase promoter, pUC origin, phage f1 origin of replication, *lacZα*, $Amp^R$ , $Kan^R$ | Qiagen |
| pJET 1.2/blunt | rep (pMB1),T7 RNA-Polyerase promoter, modified $P_{lac}$ promoter for expression of *eco47IR* and positive selection (PlacUV5), $Amp^R$ | Fermentas |
| pK18 | pUC-derived, *LacZ*' α-complementation system, (KanR) + *oriT* | Pridmore et al., 1987 |
| ΔpK18oriT*fkbL*apra | pk18-Derivate with the inactivation construct for *fkbL* | This work |
| pYT9 | pJOE2775-Derivate, $Amp^R$ | Tiffert et al., 2008 |
| pET30 Ek/LIC | Ligation Independent Cloning (LIC), T7 promoter, T7 start of transcription, phage f1 origin of replication, N-terminal His-tag and S-tag, C-terminal His-tag, T7 terminator, *lacI* coding sequence, pBR322 *ori*, $Kan^R$ | Novagen |
| pET30/*pip* | Derivate pET30 with *pip* (from *A. friuliensis*) | This work |
| pET30/*pip*$^{*\ E60A}$ | Derivate pET30 with *pip*$^{*\ E60A}$ | This work |
| pET30/*pip*$^{*\ E60Q}$ | Derivate pET30 with *pip*$^{*\ E60Q}$ | This work |

| | | |
|---|---|---|
| pET30/*pip*$^{*\ E60L}$ | Derivate pET30 with *pip*$^{*\ E60L}$ | This work |
| pET30/*pip*$^{*\ I91V}$ | Derivate pET30 with *pip*$^{*\ I91V}$ | This work |
| pET30/*pip*$^{*\ D233N}$ | Derivate pET30 with *pip*$^{*\ D233N}$ | This work |
| pET30/*pip*$^{*\ V58L}$ | Derivate pET30 with *pip*$^{*\ V58L}$ | This work |
| pET30/*pip*$^{*\ V58A}$ | Derivate pET30 with *pip*$^{*\ V58A}$ | This work |

Suppl. Tab 2. Strains and plasmids used in this study.

| Oligonucleotides | Sequences 5´-3´ | Reference |
|---|---|---|
| pipAfwNdeI | CATATGATGGAGACCTGGGTCCTGG | This work |
| pipArevBglII | AGATCTTCAGTGGGCGGGGGC | This work |
| pipfwNdeI | CATATGATGGATACGCTCCTGCTGAC | This work |
| piprevEcoRI | GAATTCGGTCAGCTGTAGGGGTTGAG | This work |
| fkbLEX_fwNdeI | CATATGATGCAGACCAAGATCCTGCGTG | This work |
| fkbLEX_revBglII | AGATCTTCACCACGGCAGCGAGTAGG | This work |
| fkbPEX_NdeI | CATATGGTGACACCGGACGGCAAGAG | This work |
| fkbPEX_neuHindIII | AAGCTTCTACTCGCTTCCCACGG | This work |
| fkbLup_fw | TCTAGAGCCGGGAGGGCCAGCGC | This work |
| fkbLup_rew | CATATGGTGGTGACGCCGGCCGGG | This work |
| fkbLdown_neu_fw | TGATATCGAGCGTCGTGGTGGTG | This work |
| fkbLdown_neu_rew | AAGCTTCGGCGCAACACTCGATAC | This work |
| Apranewfw | TAACATATGGGAGGCCAAACGGCATTG | This work |
| Apranewrev | ACAGATATCGGCCCACAGAATGATGTCAC | This work |

| | | |
|---|---|---|
| oriT_ A | GCTAGCGGCCTCCGACTAACGAAAAT | This work |
| oriT_B | GCTAGCTCTTTTCCGCTGCATAACCC | This work |
| dapAfw_NdeI | CATATGATGGCTCCGATCCCCACTC | This work |
| dapArev_BglII | AGATCTTCAGAGCTGGACGCCTCCG | This work |
| Kanfw_BlpI | AAGCTCAGCGCTTCACGCTGCCGCAAGCACTCA | This work |
| NeurevKanBlpI | TGCTGAGCAGGGGTGGGCGAAGAACTCCAGCAT | This work |
| lysC_fw_NdeI | CATATGGTGGGCCTTGTCGTGCAG | This work |
| lysC_rev_HindIII | AAGCTTTCATCGGCCGGTGCCTC | This work |
| LysCTyrfw | GAATGTGTACGCGGCCACCACCGCTCTGACCGAC | This work |
| LysCTyrrew | GGTGGCCGCGTACACATTCTGGACGATCATGTCCAG | This work |
| erme_lysC_Bam1 | GGATCCTTAGCGTCCGGTGCCTG | This work |
| erme_lysC_Bam2 | GGATCCCGCGTTGGCCGATTC | This work |
| pippETfw | GACGACGACAAGATGGATACGCTCCTGCTGAC | This work |
| PippETrew | GAGGAGAAGCCCGGTCAGCTGTAGGGGTTGAG | This work |
| Glu60Glnfw | GCGTCATCCAGTGGATGCCGCACC | This work |
| Glu60Glnrew | CATCCACTGGATGACGCCGGTGTCAC | This work |
| Glu60Leufw | GCGTCATCCTGTGGATGCCGCACC | This work |
| Glu60Leurew | CATCCACAGGATGACGCCGGTGTCAC | This work |
| Glu60Alafw | GCGTCATCGCCTGGATGCCGCACC | This work |

| | | |
|---|---|---|
| Glu60Alarew | CATCCAGGCGATGACGCCGGTGTCAC | This work |
| Asp233Asnfw | CGGCGCCAACCTCGTCGGCAAGTTCG | This work |
| Asp233Asnrew | GACGAGGTTGGCGCCGATCGCGTTG | This work |
| Val58Leufw | GACACCGGCCTGATCGAGTGGATGC | This work |
| Val58Leurew | CACTCGATCAGGCCGGTGTCACCGGG | This work |
| Val58Alafw | GACACCGGCGCCATCGAGTGGATGC | This work |
| Val58Alarew | CACTCGATGGCGCCGGTGTCACCGGG | This work |
| Ile91Valfw | CCGACGGTCATCGGCACGCTGACC | This work |
| Ile91Valrew | CCGATGACCGTCGGCAGGTTGAGG | This work |

Suppl. Tab. 3. Oligonucleotides used in this study
